# A Multi-Domain Task Battery Reveals Functional Boundaries in the Human Cerebellum

**DOI:** 10.1101/423509

**Authors:** Maedbh King, Carlos R. Hernandez-Castillo, Russell A. Poldrack, Richard B. Ivry, Jörn Diedrichsen

**Affiliations:** Department of Psychology, University of California, Berkeley, USA; Brain and Mind Institute, Western University, London, Ontario, Canada; Department of Psychology, Stanford University, Stanford, USA; Department of Statistical and Actuarial Sciences, Western University, London, Ontario, Canada; Department of Computer Science, Western University, London, Ontario, Canada

**Keywords:** Cerebellum, Task-Based fMRI, Functional Parcellation, Functional Topography, Multi-Domain Task Battery (MDTB

## Abstract

There is compelling evidence that the human cerebellum is engaged in a wide array of motor and cognitive tasks. A fundamental question centers on whether the cerebellum is organized into distinct functional sub-regions. To address this question, we employed a rich task battery, designed to tap into a broad range of cognitive processes. During four functional magnetic resonance imaging (fMRI) sessions, participants performed a battery of 26 diverse tasks comprising 47 unique conditions. Using the data from this multi-domain task battery (MDTB), we derived a comprehensive functional parcellation of the cerebellar cortex and evaluated it by predicting functional boundaries in a novel set of tasks. The new parcellation successfully identified distinct functional sub-regions, providing significant improvements over existing parcellations derived from task-free data. Lobular boundaries, commonly used to summarize functional data, did not coincide with functional subdivisions. This multi-domain task approach offers novel insights into the functional heterogeneity of the cerebellar cortex.

## Introduction

Converging lines of research provide compelling evidence that the cerebellum is engaged in a broad range of cognitive functions, well beyond its historical association with sensorimotor control and learning^1^. Anatomical tracing studies in nonhuman primates have revealed reciprocal connections with parietal and prefrontal association cortices^2^. Individuals with lesions to the cerebellum exhibit behavioral impairments on tasks designed to assess non-motor processes such as duration discrimination, attentional control, spatial cognition, emotion perception, and executive and language function. Perhaps most intriguing, neuroimaging studies consistently reveal activations of the cerebellar cortex during a diverse set of motor, cognitive, and social/affective tasks^3^.

This raises the question of whether the cerebellum can be meaningfully subdivided into a discrete set of regions, reflecting distinct functional contributions across diverse task domains. In contrast to the cerebral cortex, the cytoarchitectonic organization is remarkably uniform across the entire cerebellar cortex. Due to this homogeneity, neuroimaging and neuropsychological studies have mostly relied on the macroanatomical folding of the cerebellum along the superior to inferior axis into 10 lobules (numbered I-X)^4^. More recently, functional parcellations based on task-free fMRI data have been proposed^5–7^. However, the degree to which these proposed boundaries correspond to functional divisions remains unclear. Task-based studies have been limited by the lack of a comprehensive neuroimaging data set. A few studies have employed data sets involving multiple tasks^7, 8^, but the small number of task conditions (<7) and the lack of a common measurement baseline have made it difficult to derive and evaluate a comprehensive task-based functional parcellation. The functional heterogeneity of the cerebellum has also been explored using meta-analytic approaches^9^, which have the disadvantage that they require data sets to be combined across different groups of participants.

In the present study, we aimed to fully characterize the functional organization of the cerebellar cortex by employing two large and diverse task sets comprising 47 unique conditions, designed to engage a broad range of sensorimotor, cognitive, and social/affective processes. Using a block design, activation for each task was measured over four fMRI scanning sessions against a common baseline. Our task set was successful in eliciting activation across the entirety of the cerebellar cortex, allowing us to derive a novel parcellation in which we characterized the functional profile of cerebellar sub-regions in unprecedented detail. The breadth of the task sets also enabled us to summarize the functional specialization of the regions in terms of the underlying latent motor, cognitive, and social/affective features.

We developed a novel metric to evaluate the strength of the proposed functional boundaries. This allowed us to address a fundamental question; namely whether there *are* distinct functional regions in the cerebellum, or whether the functional specialization is better described in terms of continuous gradients^7^. The approach is predicated on the idea that if a boundary between two regions divides functionally heterogeneous regions, then the activation pattern for two voxels that lie within the same region should be more correlated than voxel pairs that span a boundary. The metric takes into account the fact that the functional similarity of two voxels will depend on their spatial distance. Critically, a meaningful functional parcellation needs to be predictive of boundaries for the activation patterns elicited by a different set of tasks. Using this approach, we demonstrate that the cerebellum has discrete functional regions, and that our MDTB parcellation is superior to alternatives in predicting functional boundaries. The new functional parcellation of the cerebellar cortex provides an important step towards understanding the role of the cerebellum across diverse functional domains.

## Results

To obtain a comprehensive functional parcellation of the cerebellar cortex, we developed a multi-domain task battery (MDTB) of 26 tasks comprising 47 unique task conditions (Fig 1a; Supplementary Table 1), selected to encompass a wide range of processes required for motor, cognitive, and affective/social function. To avoid strong learning-related changes, 24 healthy individuals completed extensive training on the task protocol (∼14 h) before scanning. During scanning, each task was performed once per imaging run for a 35 second block (Fig 1b). This task design ensured that all tasks were measured against a common baseline, allowing for between-task comparison. To make this approach feasible, the tasks were split into two sets (Fig 1a), and each task set was tested in two separate fMRI scanning sessions, resulting in a total of ∼5.5 hours (19,136 timepoints) of functional data per participant.

**Figure 1.**
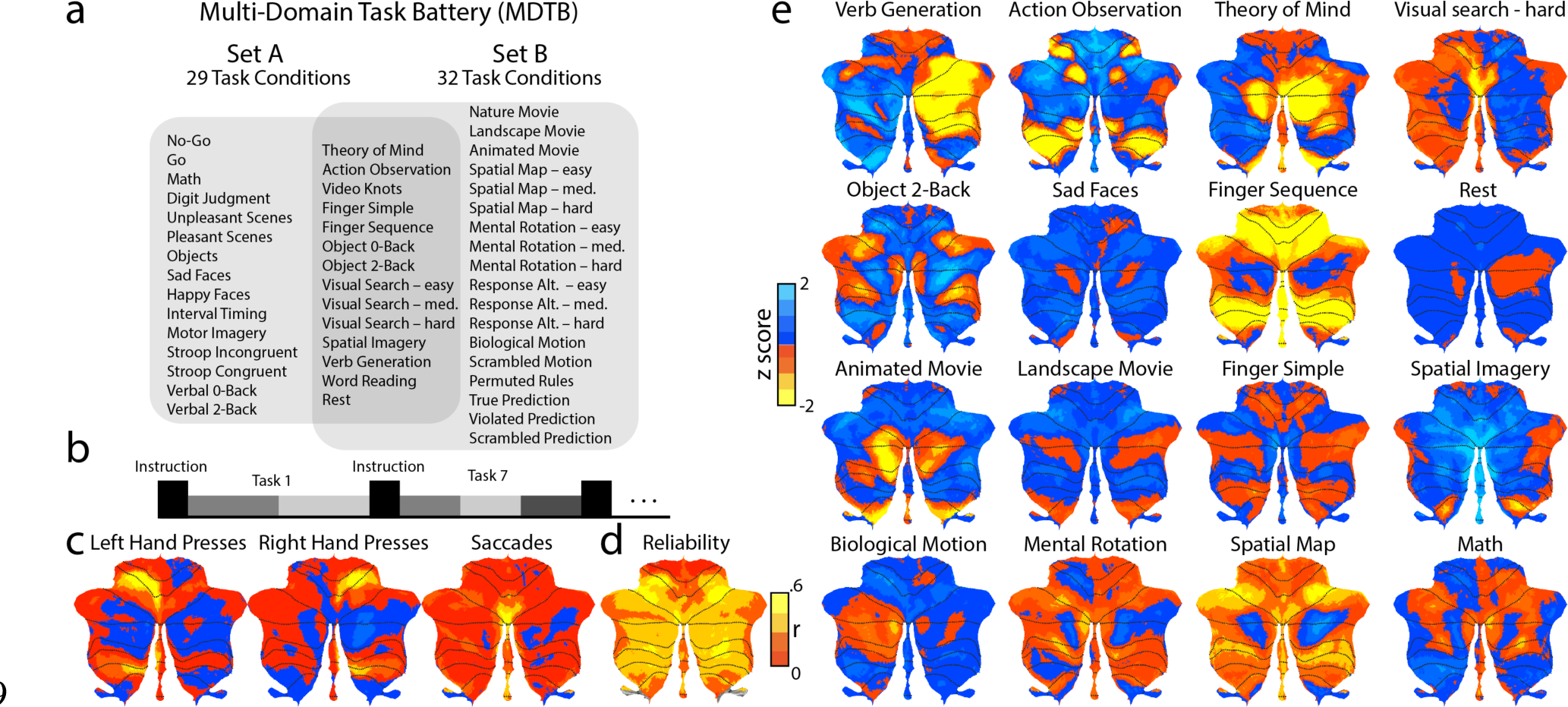
(**a**) Experimental Design. A total of 4 fMRI scanning sessions were collected on the same set of participants, using 2 tasks sets. Each set consisted of 17 tasks, with 8 tasks in common. The tasks were modeled as 29 task conditions in set A, and as 32 in set B, with 14 task conditions common across both task sets. (**b**) Timing of each task: 5 s instruction period followed by 30 s of task execution. Tasks consisted of a different number of task conditions (gray bars, range 1-3). (**c**) Unthresholded, group-averaged motor feature maps, displayed on a surface-based representation of the cerebellar cortex^10^. (**d**) Across-session reliability (r) of activation patterns for each voxel. (**e**) Group activation maps for selected tasks, corrected for motor features. Red-to-yellow colors indicate higher levels of activation and blue colors denote decreases in activation (relative to the mean activation across all conditions).

### Identification of motor features from the multi-domain task battery

As a first step, we sought to identify regions within the cerebellum in which the hemodynamic response was closely tied to motor function, specifically hand and eye movements. Our experimental design did not include specific contrasts that isolated each motor component. Instead, we varied the motor demands across task conditions; for example, the finger sequencing task involved ∼40 left and right finger responses per 30 s, the theory of mind tasks entailed two left hand responses, and the movie tasks had no overt response. We then used a feature-based approach to identify movement-related activation patterns. The motor feature model included the number of left and right hand responses for each task as measured in the scanning sessions, and the mean number of saccadic eye movements, measured in the final training session outside of the scanner. Using regularized regression (see methods), we could estimate the activation across tasks attributable to motor involvement.

Left and right hand movements were associated with activation increases in the two hand motor areas of the cerebellum (Fig 1c), the anterior hand region located on the boundary of lobules V and VI, and the inferior region in lobules VIIIb^11^. Saccadic eye movements elicited activation in the posterior vermis (especially vermis VI), consistent with the location of the oculomotor vermis in the macaque monkey^12^. Compared to previous contrast-based human fMRI studies^13^, which have yielded relatively inconsistent results, our feature-based mapping approach resulted in an extraordinarily clear localization of eye-movement activation to the oculomotor vermis. While these results mainly confirm the well-known functional localization within the cerebellum for movement, they demonstrate that a broad task-based approach without tightly matched control conditions provides a powerful means of revealing functional organization.

### Multi-domain task battery elicits varied activation patterns across the cerebellum

We then aimed to characterize activation patterns invoked by the task conditions that could not be explained by the basic motor features. Overall, our task sets were able to elicit strong and distinguishable patterns of activation (Fig 1e; see Fig S1 for the full set of group maps) across the cerebellar cortex. To determine the reliability of the activation patterns, we calculated the correlation of the individual, unsmoothed task-activation profiles for each voxel across the two sessions of each set. On average, these task activation profiles were reliable (set A: r=.43, CI: .39-.46; set B: r=.42, CI: .37-.46; see Fig S2 for individual participant maps). The resulting voxel-wise reliability map (Fig 1d) confirmed that this was the case for the entire cerebellar cortex, with the exception of lobules I-IV. These lobules are associated with foot movements^11^ and postural responses^14^, features that are absent in our task sets.

Qualitatively, the activation patterns elicited by our task sets replicated numerous results obtained in previous neuroimaging studies that had focused on a single task domain. For example, highly right-lateralized activation throughout Crus I, Crus II, and VIIb was observed with the verb generation task^8^ while left-lateralized activation throughout Crus, I, Crus II was demonstrated with the biological motion task. Similarly, the verbal and picture N-Back tasks activated two distinct lateral regions of lobules VII, in agreement with previous working memory studies^10^. Recent evidence for activation of medial Crus I and Crus II by movie tasks was also corroborated here^59^.

The task-activation maps also demonstrated some new insights, which have not been (or not as clearly) reported in the previous literature. The rest condition (contrasted against the mean of all the other conditions) was associated with bilateral activation in a mid-hemispheric region in Crus I and II, effectively forming the cerebellar component of the default-mode network^5^. Similar cerebellar regions were strongly activated during the theory of mind task^10^ and the movie tasks^59^. The finger sequencing and visual search tasks led to strong activation in cerebellar hand and eye-movement related areas, respectively. Given that these activation maps were corrected for activation explainable by the sheer number of hand and eye movements, the maps indicate that these areas are especially activated during complex movements or conditions with high attentional demands. Finally, the action observation task elicited activation in a distinct set of areas surrounding the motor areas of the cerebellum, especially in the posterior motor representation. When using a dissimilarity measure to construct a representational space for all tasks (Fig S4a), the action observation condition can be seen to produce one of the most unique activity patterns.

The passive picture viewing tasks (i.e. sad faces) did not elicit much activation in the cerebellum. This is generally consistent with the notion that the cerebellum does not receive cortico-pontine projections from the inferior temporal cortex, a pathway involved in the recognition of visual scenes and objects. To quantify this observation, we tested the activation patterns of all possible task conditions against each other. While over 95% of the pairwise comparisons were significant (uncorrected p<.001 level), the most notable exceptions were pairs of the picture-viewing tasks (Fig S4a). In contrast, passively watching engaging movie snippets (Nature movie, Animated movie) resulted in reliable and specific activity patterns (Fig 1e, Fig S1), likely related to processes required for action perception and social cognition.

### Cerebellar lobules do not reflect functional subdivisions

One way to summarize these activation patterns is to subdivide the cerebellum into functionally distinct regions. This approach, however, is only meaningful if there are stable functional subdivisions in the cerebellum that generalize across tasks. To address this fundamental question, we developed a new evaluation metric, which we refer to as the distance controlled boundary coefficient (DCBC). The metric is predicated on the idea that if a boundary divides two functionally heterogeneous regions, then any equidistant pair of voxels that lie within a region should have activation profiles that are more correlated with each other than two voxels that are separated by the boundary (Fig 2a, see methods). Specifically, we calculated correlations between voxel pairs using a range of spatial bins (4 mm to 35 mm), and then used the difference between the mean correlation of within-region pairs and between-region pairs as our criterion. This method extends standard clustering assessment tools (such as the silhouette coefficient) to account for spatial distance.

**Figure 2.**
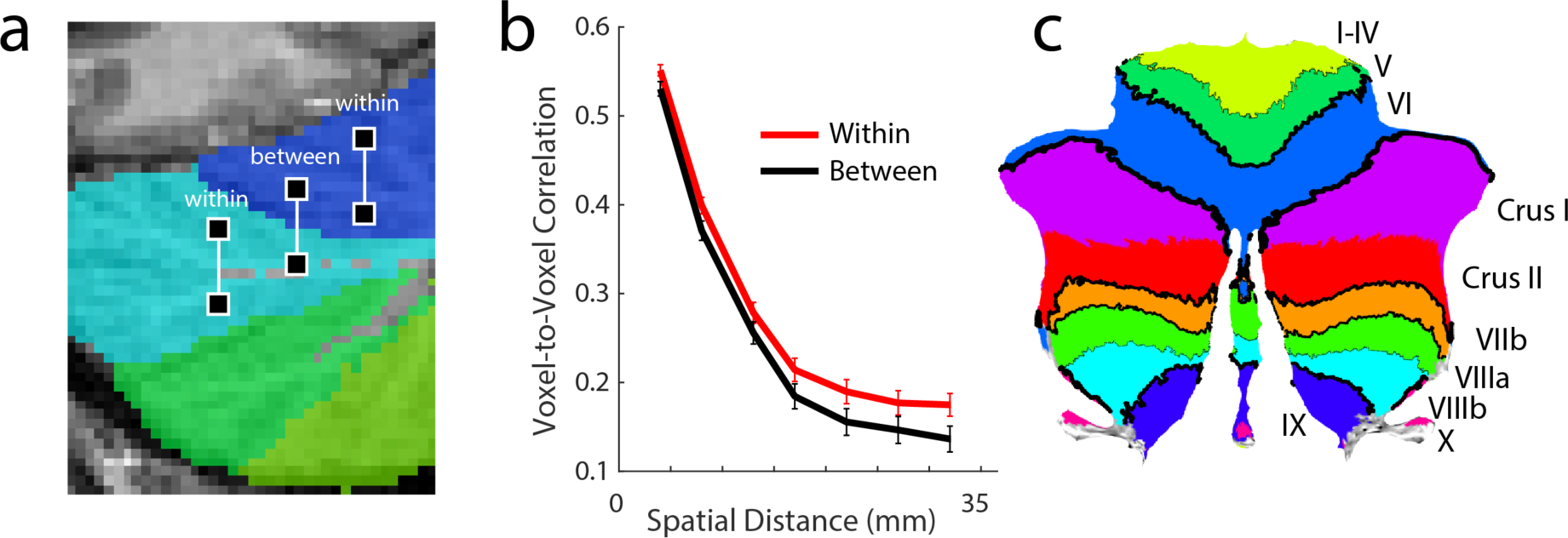
Distance-corrected boundary coefficient (DCBC). **(a**) Correlations between all pairs of voxels with the same distance were calculated and averaged depending on whether they were “within” or “between” regions. Voxel-pairs were then binned according to spatial distance in the volume (4-35mm in steps of 5mm). (**b**) Correlation as a function of spatial distance for lobular boundaries. The DCBC is defined as the difference in correlation (within-between) within each distance bin. (**c**) Strength of the boundaries for the lobular parcellation, with the thickness of the black lines indicating the DCBC value.

We first employed this evaluation method to determine the degree to which functional boundaries follow the major lobular subdivisions. While we would not expect a 1:1 correspondence between lobular and functional boundaries^5^, this is a question of high practical importance given that lobular boundaries are commonly used to define regions-of-interest for interpreting functional activations in the cerebellum. Notably, the correlation between voxels within a lobule was not much greater than the correlation between voxels that spanned a lobular boundary (Fig 2b). The correlations, averaged over distances of 4 mm to 35 mm, were r=0.28 (0.26 – 0.30) within-lobules and r=0.25 (0.05 – 0.46) between-lobules (95% confidence interval). While the difference was significant (t_23_= 4.703, p<.01), the size of the difference was small (DCBC=0.03). Thus, lobular boundaries do not reflect strong functional subdivisions in the cerebellum.

As well as providing a global evaluation criterion, the DCBC can also be used to evaluate the strength of individual boundaries (Fig 2c). For example, the superior posterior fissure separating lobule VI from VII was the strongest lobular boundary (DCBC=.127), while the primary fissure, which serves as first principle subdivision of the cerebellum, was relatively weak (DCBC=.069). The boundary separating Crus I and Crus II did not predict any functional specialization (DCBC=0). In sum, a lobular parcellation poorly identifies functional regions within the cerebellar cortex.

### MDTB parcellation uncovers strong functional boundaries

Next, we asked whether the MDTB parcellation would more clearly identify functional boundaries in the cerebellum. While functional parcellations will invariably yield boundaries from a given task set, it cannot be assumed that these boundaries will generalize to new tasks. We first estimated a group-based parcellation using all of the MDTB data. Using convex semi-nonnegative vector factorization^15^, we decomposed the N (tasks) x P (voxels) data matrix into a product of an N x Q (regions) matrix of task profiles and a Q x P matrix of voxel weights. The voxel weights, but not the task profiles, were constrained to be non-negative. Using a winner-take-all approach, we then assigned each voxel to the region with the highest weight. Fig 3b shows the resulting parcellation using 10 regions.

**Figure 3.**
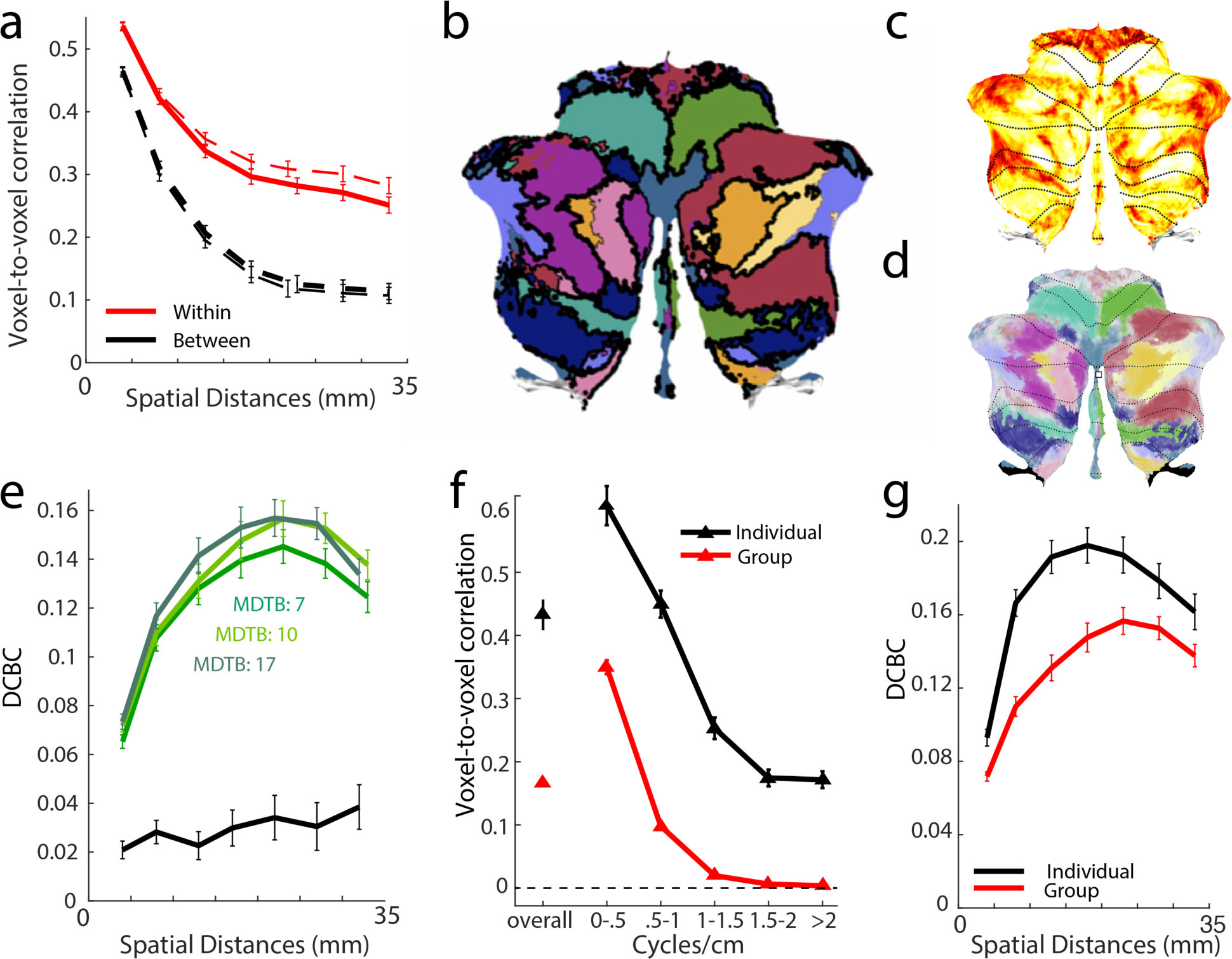
The MDTB parcellation reveals functional boundaries in the cerebellar cortex. **(a)** Correlations for “within” (red) and “between” (black) voxel-pairs for the MDTB parcellation (10 regions). Solid lines indicate the values for the cross-validated estimates; the dashed lines are the estimates for the full parcellation. **(b)** 10-region MDTB parcellation. The DCBC for each boundary is visualized by the thickness of the black lines. **(c)** Proportion of samples in the bootstrapped analysis (participants) in which the voxel was assigned to the same compartment as in the original parcellation. Most voxels had an assignment certainty >60%. **(d)** Visualization of boundary uncertainty, using the color scheme in panel b), but adjusted such that the degree of transparency is indicative of the uncertainty of the assignment. Voxels that were assigned to a single compartment on less than 50% of the cases are shown in gray **(e)** DCBC as a function of the spatial distance for the lower bound of the three MDTB parcellations (colored lines) and lobular parcellation (black line). (**f**) Within-subject (black) and between-subject (red) reliability of activation patterns overall, and across different spatial frequencies. (**g**) DCBC as a function of voxel distance for the 10-region group parcellation (red) and average of the 10-region individual parcellations (black). Only shown are the cross-validated estimates of the prediction performance for novel tasks.

For this parcellation, the average DCBC was .157 (Fig 3a, dashed line, t_23_=34.84, p<1e-10), a value higher than that obtained for the strongest lobular boundary. However, because the training and evaluation data overlap, this evaluation will be biased in favor of the MDTB parcellation. For a more conservative estimate, we determined the parcellation based on all task conditions from set A, and evaluated the boundaries using the unique tasks from set B. We repeated this out-of-sample generalization test in the other direction and averaged the two values. Using this approach, the average DCBC was .130, only slightly lower that the non-crossvalidated estimate (Fig 3a; t_23_=24.056, p<1e-10). The cross-validated DCBC will underestimate the true predictive power of the full parcellation, with true performance on a novel task likely falling between the cross-validated and non-crossvalidated DCBC. To remain conservative, we only report the cross-validated DCBC estimates for the remainder of the paper.

The exact form of a parcellation depends on the specified number of regions. We also derived a 7-region parcellation (Fig S5d) and a 17-region parcellation (Fig S5f). While the 7-region parcellation performed slightly poorer than the 10-region parcellation (DCBC=.121, t_23_=-3.901, p=.00072), the performance of the 17-region and 10-region parcellations was comparable (DCBC=.133, t_23_=1.55, p =.133). Quantitatively, there was reasonable agreement across the different MDTB parcellations (Fig S5h, Fig 4f). Visually, however, there were some marked differences in the functional subdivisions for the different parcellations. While our results clearly show that the MDTB parcellations reflect true functional boundaries in the cerebellum, they also make clear that there are a number of equivalent ways to subdivide the cerebellum. Thus, the exact choice of a “final” parcellation is constrained by practical considerations. Here, we have chosen to focus on the 10-region parcellation, as it provides a useful level of resolution for a full functional characterization.

**Figure 4.**
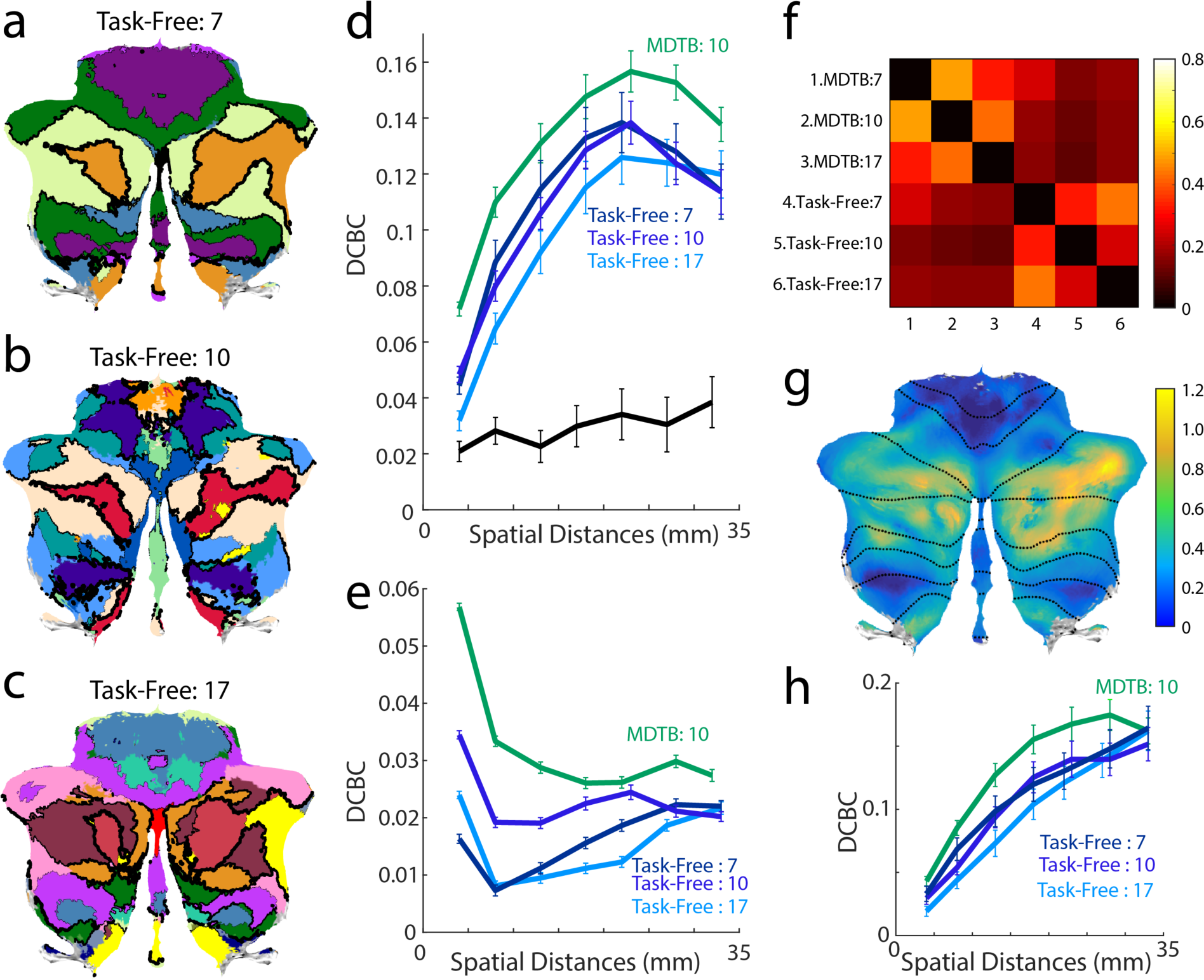
Comparison of MDTB and task-free parcellations of the cerebellum. **(a-c)** Cerebellar parcellations based on task-free data with 7, 10, and 17 regions^6^. Thickness of the black lines indicates the DCBC for the corresponding boundary. **(d)** DCBC for different spatial distances for lobular (black), task-free (dark-light blue) and MDTB (green) parcellations. The MDTB parcellation was evaluated in a cross-validated fashion (see text). (**e**) Evaluation of the same parcellations on task-based data from 186 HCP participants**. (f**) Matrix of adjusted Rand coefficients (AR) between three versions of the MDTB parcellations (7, 10, 17) and the three task-free parcellations (7, 10, 17). (**g**) Correspondence between MDTB and task-free parcellations. A value of 1 (yellow-green) indicates that the AR between task-free and MDTB parcellations is the same size as the AR between MDTB parcellations. Lower values (blue-green) indicate weaker agreement between task-free and MDTB parcellations compared to MDTB by itself. (**h**). Evaluation of the MDTB parcellation (derived only from Set A) and the task-free parcellations on the three movie tasks from Set B.

To assess the stability of the parcellation, we conducted a bootstrap analysis, both across participants and task conditions (for details, see methods). The mean Rand coefficients between each of the new parcellations and the original parcellation was 0.628 (95% CI=0.54-0.71) and 0.632 (95% CI=0.54-0.72) for the bootstrap analyses across participants and task conditions, respectively. To quantify the uncertainty of specific boundaries, we calculated the proportion of bootstrap samples for which each voxel was assigned to the same compartment as in the original parcellation (Fig 3c). Overall the consistency was very good for most of the cerebellum (Fig 3d). Areas of high uncertainty were lobules I-IV, likely a consequence of the fact that we did not include foot movements in our task-battery.

The parcellations described above were based on group data; but it is important to also consider the variability in functional organization across individuals. To quantify inter-subject variability, we compared the correlation between the task-activation maps across participants to the within-subject reliability across the two sessions for each set (Fig 3f). Overall, ∼36% of the pattern variance was shared between individuals, whereas ∼59% reflected idiosyncratic patterns. A spatial-frequency decomposition of the patterns (see methods) revealed that commonalities across participants were restricted to the low spatial frequencies (< 1 cycle/cm; activations of more than 5mm in size), while the fine-grained patterns were purely idiosyncratic to the participant. Indeed, a parcellation derived from the functional data from the individual significantly outperformed the group parcellation in predicting functional boundaries on that same individual (Fig 3g; t_23_=5.883, p=1e-5).

In summary, using the MDTB data, we were able for the first time to quantitatively demonstrate the existence of distinct functional regions in the human cerebellum. Our results clearly advocate the adoption of a functional parcellation, to replace lobular subdivisions as a tool to summarize functional cerebellar data.

### Task-free parcellations identify overlapping, but weaker boundaries

Prior work has leveraged the correlational structure of task-free (or “resting state”) fMRI data to derive various parcellations of the cerebellum, with published reports using 7^5^, 10^6^, or 17^5^ regions (Fig 4a-c). These parcellations are only moderately consistent with each other (Fig 4f), with an average adjusted Rand index (AR) of 0.33 (0 = no communality; 1 = perfect match). Correspondence between the different MDTB parcellations was slightly higher (AR=0.40), indicating more stability across the MDTB parcellations. The average AR between the MDTB and task-free parcellations was 0.16, underscoring that there are systematic differences between the two approaches. To explore regions in which task-free and MDTB parcellations diverge, we conducted a searchlight analysis, computing the Rand coefficient locally using a 1cm-radius sphere for each pair of parcellations. The results showed that the correspondence between task-free and MDTB parcellations is best in the mid-lateral areas of lobule VII around the “default-mode” network. In this region, the agreement between the MDTB and task-free maps (Fig 4g) was similar to the agreement between the MDTB maps themselves (Fig S5h). In contrast, in more lateral aspects of lobule VII, and especially areas engaged in motor control or action observation, the correspondence between task-free and MDTB parcellations was much weaker. This is likely due to the relatively low consistency among the task-free parcellations (Fig S5g).

We evaluated how these task-free parcellations were able to predict functional boundaries in our MDTB data. For all task-free parcellations, the within-region correlations were significantly higher than the between-region correlations (t_23_ =12.929, p<1e-10). The average DCBC for the 7, 10, and 17-region parcellations was .109, .105, and .096 respectively, substantially higher than the lobular parcellation (t_23_=16.388, p<1e-10; Fig 4d). Thus, all of the task-free parcellations are, to some degree, able to predict functional boundaries. However, the average task-free DCBC was significantly lower than the “lower bound” for our MDTB 10-region parcellation (t_23_=5.844, p<1e-5). This indicates that the MDTB parcellation outperformed the task-free parcellations in predicting functional boundaries on a novel set of task conditions.

Although the evaluation of the MDTB parcellation was fully cross-validated, with no direct overlap between conditions used for training and evaluation (see also Fig S6), we also wanted to ensure that the superior performance of the MDTB parcellation would generalize to a completely separate dataset. To this end, we evaluated the MDTB and task-free parcellations using task-based data from 186 participants tested as part of the Human Connectome Project (HCP). Again, the MDTB 10-region parcellation significantly outperformed the three task-free parcellations (7-region: t_185_ =22.756, p<1e-10; 10-region: t_185_=14.36, p<1e-10; 17-region: t_185_ =27.629, p<1e-10).

To ensure that the higher predictive power of the MDTB parcellation was not solely driven by regions associated with motor control, we re-evaluated the DCBC using only the three movie tasks (Fig 4h). Even though these conditions did not demand any overt movement, the advantage of the MDTB over the 7-region (t_23_=2.41, p=.024), 10-region (t_23_=5.028, p<1e-5), and 17-region (t_23_=4.754, p<1e-5) task-free parcellation remained significant. Overall, these results demonstrate that the advantage of the MDTB over task-free parcellations extends to new data sets and to conditions that do not involve active tasks.

### Characterizing activation by cognitive features

An important advantage of a task-based approach is that we can make inferences about the processes that activate the cerebellar cortex. While our data-driven parcellation identified boundaries, such that each region had a relatively homogenous functional profile, the profiles themselves are not immediately interpretable. One way of characterizing the task profiles is to use predefined and non-orthogonal features^16^. We have already successfully applied this approach when characterizing the activation patterns elicited by motor features (Fig 1c), which could be directly operationalized as the number of finger and eye movements. To extend this approach, we needed to describe each task condition in terms of its underlying cognitive features. We therefore turned to Cognitive Atlas, an online cognitive ontology^17^, that summarizes the current consensus in cognitive science of the processes associated with a large array of tasks. To construct a feature space, each of the task conditions was rated on each of the cognitive concepts (see methods). We then estimated feature weights for each region using non-negative regression. For visualization purposes, we depicted the top three feature weights for each region (Fig 5).

**Figure 5.**
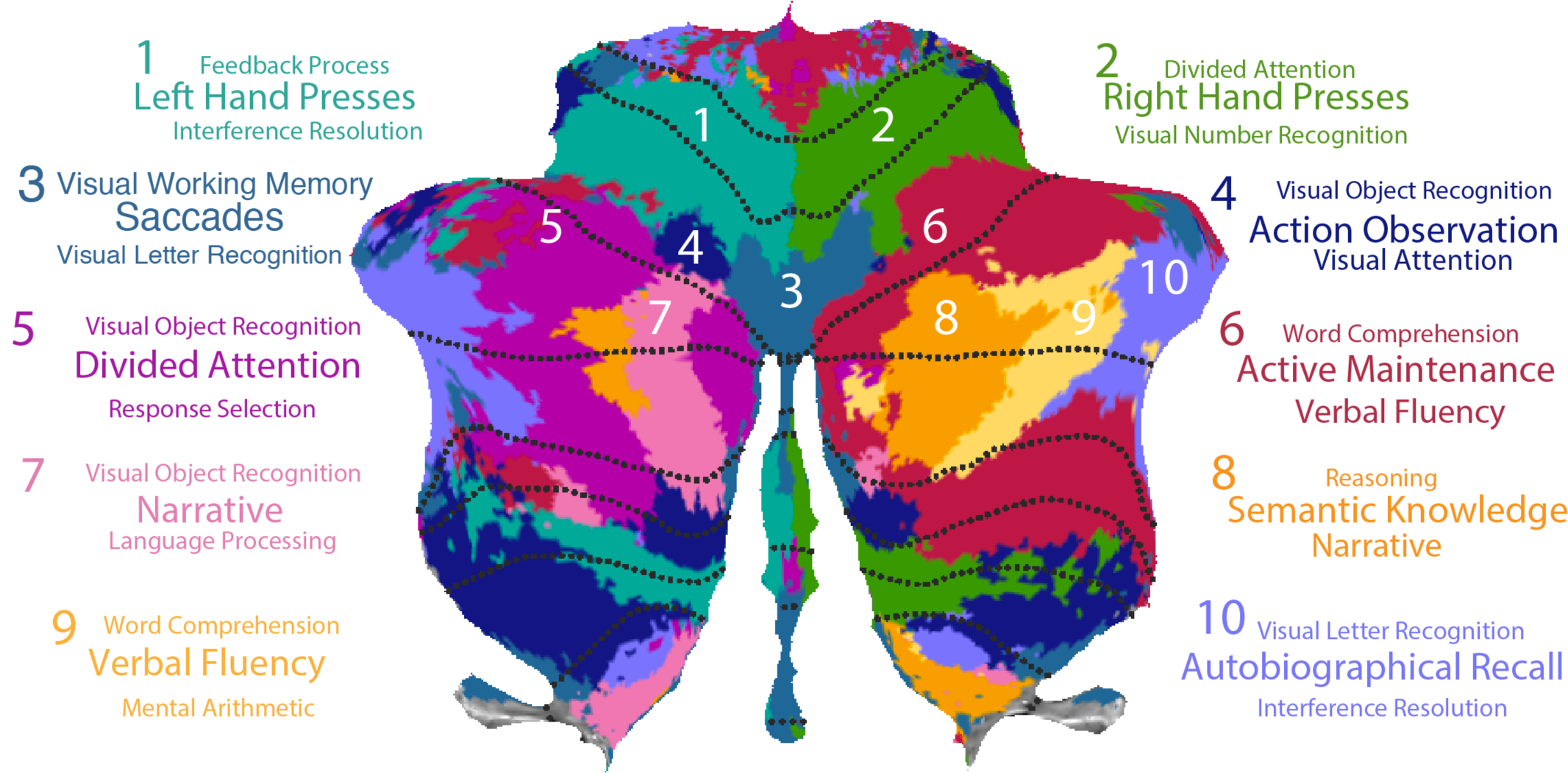
Cognitive descriptors for the 10 functional regions in the MDTB parcellation. Three features that best characterize each region are listed. Font size indicates the strength of these feature weights.

The dominant features describing the three motor regions (regions 1, 2, 3) were left-hand, right-hand and saccadic eye movements, respectively. The posterior associative motor region (region 4) was driven predominantly by action observation. For the other regions the dominant features related to a range of cognitive processes. Regions 5 and 6 in the mid-hemispheric aspects of Crus I/II, lateralized to the left and right hemisphere respectively, were associated with attention-related features such as divided attention (region 5) and active maintenance (region 6). In contrast to the task-free parcellations, the MDTB map assigned homologous areas in the left and right hemispheres to different regions, suggesting some degree of hemispheric asymmetry within the cerebellum. Nonetheless, the task-activity profiles between homologous regions were strongly correlated (Fig S7).

More medially in both hemispheres were regions 7 and 8, best described by features associated with either narrative (region 7) or semantic knowledge (region 8). Activity in right-hemispheric region 9, lateral to region 8, was best explained by features related to language processing (e.g., verbal fluency and word comprehension). Finally, region 10, encompassing the most lateral aspects Crus I/II was dominated by autobiographical recall. This region shows strong task-free correlations with the frontal pole^5^. Overall, activity in the larger proportion of the cerebellum was explained by features related to cognitive, rather than motor processes^18^.

## Discussion

### Summary

The aim of this study was to derive a comprehensive picture of the functional organization of the human cerebellum. To do this, a group of participants was scanned over the course of four fMRI sessions while performing a diverse multi-domain task battery composed of 26 unique tasks comprising 47 conditions. The task-evoked activation patterns were leveraged to derive a functional parcellation of the cerebellar cortex. We employed a new technique to quantitatively evaluate the boundaries of this parcellation in comparison to parcellations based on lobular structure and task-free fMRI data. The MDTB parcellation successfully predicted functional boundaries when tested with a novel set of tasks, outperforming alternative parcellations.

### Parcellations of the cerebellar cortex

The lobular architecture of the cerebellum has provided, both in neurophysiological and neuroimaging studies, the primary reference for defining sub-regions^4, 19^. The macroanatomical folding into ten lobules is well conserved across species^4^ and under strong genetic control^20^. While the results from electrophysiological^21^ and neuroimaging studies^7, 22^ have suggested that lobular boundaries do not demarcate functional subdivisions, we present here the first quantitative evaluation of this hypothesis. Indeed, lobular boundaries appear to constitute only very weak boundaries in terms of functional organization. The identified functional regions often spanned multiple lobules, with many of the boundaries traversing the cerebellar cortex along the parasagittal axis. The clear dissociation of anatomical and functional organization of the cerebellum, as revealed here, questions the value of summarizing functional and anatomical data in terms of lobular regions-of-interest.

As an alternative, we employed a diverse task battery to develop a parcellation that could comprehensively describe the functional organization of the cerebellar cortex. Critically, the group-based MDTB parcellation predicted functional boundaries in individual participants, as shown using out-of-sample generalization on the same data set, as well as generalization to a completely separate data set (HCP task data). These findings provide a compelling demonstration of discontinuities related to functional specialization across the cerebellar cortex. Evidence from meta-analyses has suggested the existence of “motor”, “cognitive”, and “affective” regions of the cerebellum^8^. However, it has also been suggested that functional variation across the cerebellar cortex may be best understood in terms of smooth gradients^7^, without definable boundaries. If this were the case, our DCBC measure, reflecting the ratio of the strength of within-region to between-region correlations, would have been near zero when tested on a novel task set. Instead, the values were positive, providing a rigorous demonstration of functional boundaries in the cerebellar cortex.

The DCBC measure also provides a means to quantify the strength of each boundary. For the MDTB parcellation, the strongest boundaries demarcated the motor areas in lobules V and VI, and lobule VIII. Furthermore, lobule VII was subdivided into an intermediate region (regions 2 and 4) and a set of surrounding regions (regions 3, 5 and 6), consistent with a mirror-reflected organization across the anterior-posterior axis of the human cerebellum^23, 24^.

An open question is whether the boundaries defined through our task-based approach relate systematically to anatomical features of the cerebellum identified by molecular techniques^21^. Specifically, studies investigating Adolase-C (Zebrin) expression in Purkinje cells in the rodent^25^ and primate brain^26^ have revealed a series of parasagittal zones across the cerebellar cortex. Olivo-cerebellar projections respect this zonal organization, with single climbing fiber inputs synapsing onto Purkinje cells that lie within a zone^25^. The organization of Zebrin-zones remains to be established in the human cerebellar cortex.

However, we suspect that the alignment with the organization observed here may not be very tight given that the cerebellar hemodynamic signal is primarily reflective of mossy fiber input^27^. Relative to climbing fibers, mossy fiber innervation patterns are likely to be patchier and more diffuse, potentially spanning multiple zonal regions^28^.

Boundaries identified from task-free fMRI data were also able to predict task-based discontinuities. This finding is in accord with similar analyses of the cerebral cortex, demonstrating that task-based activation patterns in the neocortex can be predicted to some degree by parcellations obtained from the spontaneous fluctuations in the fMRI signal during rest^29^. However, our MDTB parcellation outperformed alternative task-free parcellations^6, 22^ in predicting functional boundaries for completely different tasks within the MDTB and HCP datasets. While the MDTB parcellation was based on fewer participants than the other parcellations (24 vs. 1000), our data set entailed considerably longer scanning time/participant. One notable difference between the task-free parcellations and our MDTB parcellation was that the latter distinguished between left and right hand movement regions in both the anterior and posterior lobes, whereas task-free parcellations lump these together.

While our group-based map could predict functional boundaries in individual participants, the finer spatial details of the functional organization were idiosyncratic for each individual. Consistent with this, the individual parcellation outperformed the group parcellation in predicting functional boundaries in that individual. Of course, individual parcellations require data collection for each participant using at least a subset of our task battery. It is worth noting that each individual parcellation was obtained on almost 3 hours of data per participant, an amount of scanning time that is usually not feasible or practical. In future studies we aim to determine the required amount of data per participant, identify the best subset of tasks, and explore the possibility of combining individual and group data to derive an optimal parcellation.

### Novel insights about functional topography

An additional advantage of a multi-domain task-based approach for mapping the cerebellum is that we can not only identify functional boundaries, but we can also relate the activation patterns to the task requirements. For many of the tasks in our battery, the activation patterns were in accord with the results obtained in previous fMRI studies that have examined a single or limited set of task domains. Examples here include the verbal and object working memory tasks, and the theory of mind task.

By using a rich multi-domain task battery, we also identified functional regions that had not been observed or well-described in previous work. A large and extended region of the cerebellar cortex was activated during action observation (region 8), surrounding the anterior, but to a much larger extent, posterior hand motor region. Interestingly, the action observation region was also activated during complex movement as shown by the sequence production task. Taken together, these results suggest that anterior motor regions are more related to primary action execution, whereas posterior motor regions are more akin to a “premotor” area, perhaps associated more with action planning and action comprehension. Notably, lesions limited to the posterior cerebellum rarely lead to lasting symptoms of ataxia^30^.

A second example comes from our motor feature model, which revealed a region around vermis VI that was strongly associated with saccadic eye movements. This finding is consistent with neurophysiological data from non-human primates showing that this region is associated with oculomotor control^12^. However, prior neuroimaging studies of the cerebellum have proven controversial with respect to this issue: Some studies have also linked this area with eye movements^13^, but other studies have argued for a functional role of this region in more complex cognitive and/or affective processes^7^. Based on pilot work for this study and other unpublished observations, we have found it difficult to elicit any cerebellar activation with a simple saccadic eye movement task. In contrast, we observed robust activation in this area during the visual search task, even when accounting for the average number of saccades made during a 30 second block. Thus, our results offer a novel perspective on the functional role of this region, indicating that the hemodynamic signal here is driven by eye-movements performed under high attentional demands.

The identification of this oculomotor region is also of interest given that prior studies have suggested that vermal activation in lobule VI is associated with emotional processing^7^. Conditions in our battery designed to engage emotional and affective processing (e.g., static images of unpleasant and pleasant scenes, sad faces) only weakly activated this region. Given the strong association of this region with eye movements, it may be that the prior activations were more related to differences in saccadic eye movements between conditions, rather than the emotional and affective processing demands of the tasks. Further support for this hypothesis comes from task-free fMRI studies showing that the oculomotor vermis is functionally connected to visual regions of the cerebral cortex^5, 7^. An obvious challenge for future research is to explore how variation in eye movements, affective processing, and attentional demands interact in driving activation within this region.

## Conclusions

This paper presents a comprehensive multi-domain task battery for the human cerebellum, unique in its functional diversity of task conditions and amount of data per individual. The group and individual task contrast maps and the group parcellations are made available at diedrichsenlab.org/imaging/mdtb.htm. We anticipate that this resource will be useful for two important applications. First, the data, combined with our novel evaluation criterion, provides a quantitative assessment of functionally defined boundaries, something that has been absent in prior studies. Given that our acquisition parameters covered the entire brain, the methods presented here can be used to evaluate parcellations of the neocortex and other brain structures. Second, the novel MDTB parcellations provide an important tool to define functional regions of the cerebellum. Our analyses clearly show that the MDTB parcellations predict functional boundaries in a novel set of tasks better than existing task-free parcellations. Moreover, each region can be characterized by a rich functional task-profile, allowing for a characterization of the associated cognitive processes. For future research, the parcellation and the associated features can provide a useful guide in designing studies to test specific functional hypotheses, and to provide a reference for interpreting the results. The MDTB functional parcellation should also be of considerable utility for translational work, given the hypothesized involvement of cerebellar dysfunction in a range of neurological and psychiatric disorders^31^. A functionally-defined parcellation can help reveal dysfunction in specific cerebellar regions and cerebro-cerebellar circuits^32^, providing further insight into the interaction between the cerebellum and neocortex.

## Methods

### Participants

All participants gave informed consent under an experimental protocol approved by the Institutional Review Board at Western University. A total of 31 participants were scanned performing set A, and 26 of this original cohort returned at a later date to perform set B (mean break between sessions = 166 days; SD=153 days, with half returning about a year later and the other half having sessions separated by 2-3 weeks). The five participants who did not return for set B were not included in the analyses. Two additional participants were excluded from the analyses as they failed to complete all 32 scanning runs. Therefore, the final sample for the multi-domain task battery (MDTB) consisted of 24 healthy, right-handed individuals (16 females, 8 males; mean age=23.8 years old, SD=2.6) with no self-reported history of neurological or psychiatric illness. Right-handedness was confirmed by a score greater than 40 on the Edinburgh Handedness Inventory.

### Experimental tasks

The experimental tasks included in set A of the MDTB were chosen to elicit heterogeneous activation patterns in the cerebellum. We selected tasks to sample across a wide range of processing domains (cognitive, motor, affective, social), in many cases drawing on tasks that had previously been shown to engage the cerebellum. While recognizing that our selection process was somewhat arbitrary and that the tasks would differ on a number of different dimensions, our main criterion was to use a large battery that broadly sampled different functional domains. A full description of the tasks, along with the accompanying references is provided in Supplementary Table 1.

The design of set B was guided by the results obtained with set A. Set B included 8 tasks that had been included in set A (shared tasks, e.g., i.e. theory of mind, finger sequence) and 9 unique tasks. The shared tasks provided a means to establish a common baseline across the two task sets. This enables between-task comparison across task sets, which is done by subtracting the mean activation pattern of the shared tasks from each task set. Only tasks that were successful at eliciting activation in the cerebellar cortex in set A were included as shared tasks in set B. For example, the picture-based tasks did not elicit much activation and were omitted from set B. For some of the novel tasks, we selected conditions that are thought to assay similar processing domains as in task set A. For example, both sets included working memory tasks, but the tasks involved different stimulus dimensions (e.g., verbal working memory in set A and spatial mapping in set B). Other tasks (for example the naturalistic movie-viewing tasks) were novel in task set B.

### Experimental design

Each set consisted of 17 tasks. In every imaging run, each task was performed once for 35 s. The 35 s block was divided into a 5 s instruction period, where the task name (e.g., ‘Theory of Mind Task’), the response effector (‘Use your LEFT hand’) and the button-to-response assignment (‘1 = false belief. 2 = true belief’) were presented on the screen. This was followed by a 30 s period of continuous task performance. In general, novel stimuli were introduced across imaging runs to prevent participants from learning specific stimulus-response associations. The one exception was the motor imagery task in which participants were required to imagine playing a game of tennis. The number of trials within the 30 s block varied from 1 (e.g., the movie viewing and mentalizing tasks) to 30 (e.g., Go-No-Go task). Most tasks involved 10-15 trials per block. The motivation for testing all tasks within a scanning run, as opposed to testing one task in each run, was to ensure a common baseline for all tasks, enabling between-task comparisons.

Three of the shared tasks (object N-Back, visual search, semantic retrieval) had a rapid, discrete trial structure (15/block), whereby each unique stimulus (picture, letter, noun) was presented for 1.6 s, with the response required to be completed within this window, followed by an inter-trial interval (ITI) of 400 ms. Three of the shared tasks had a slower discrete trial structure: sequence motor task (trials=8; stim/resp duration=4.6 s; ITI=400 ms), theory of mind (trials=2; stim/resp = 9.6/4.6 s; ITI=400 ms) and action observation (trials=2; stim duration=14 s; ITI=1 s). The remaining two shared tasks, spatial imagery and rest did not have a discrete trial structure (trials=1; duration=30 s).

Of the nine unique tasks in set A, six (interval timing, IAPS affective, IAPS emotional, verbal N-back, motor imagery, stroop, go/no-go, math, passive viewing) had the rapid discrete trial structure (1.6 s/trial and 400 ms ITI). The go/no-go task also had a rapid discrete structure, but the rate was increased to maintain a high level of attention/arousal (trials=30; stim/resp duration=800 ms; ITI=200 ms). The math task was comprised of 10 trials (stim/resp duration=2.6 s; ITI=400 ms). The motor imagery task did not have a discrete trial structure (duration=30 s).

Of the nine unique tasks in set B, six had a discrete trial structure: The prediction, spatial map, and response alternatives tasks entailed 6 trials/block (stim/resp=4.8 s; ITI=200 ms), the mental rotation task 9 trials/block (stim/resp duration=3 s; ITI=300 ms), the biological motion task 10 trials/block (stim/resp duration=3; ITI=0), and the permuted rules task task 4 trials/block (stim/resp duration=7.3 s; ITI=200 ms). The three movie-viewing tasks (landscape, animated, and nature) did not have a discrete trial structure (duration=30 s).

### Hand assignment across tasks

For each task requiring responses, the responses were made with either the left, right, or both hands using four-key button boxes. Hand assignment was consistent for sets A and B for the shared tasks. For tasks requiring 2-choice discrimination, responses were made with the index or middle finger of the assigned hand while responses for tasks requiring 4-choice discrimination were made with the index and middle fingers of both hands. By including a motor feature model in our analysis (see below), we were able to account for the motor requirements across the tasks.

### Behavioral training

For each task set, participants completed three days of training prior to the first scanning session. Training included all of the tasks with the exception of the rest condition and the three movies (set B). For each set, the three training sessions took place over the course of four to seven days (set A: mean number of days=5.2, SD=3.5; set B: mean number of days=4.4, SD=1.8).

The first day was used to familiarize the participants with the requirements for each of the 17 tasks. The participants were instructed to carefully read the instructions. When ready, they initiated a 35 s training block. The number of training blocks differed depending on the perceived level of difficulty of the task. For example, the 2AFC picture-based tasks (IAPS affective, IAPS emotional) were practiced for three blocks, while the Stroop task was practiced for seven blocks. During this training session, a run consisted of consecutive blocks of the same task. On-line feedback was provided for response-dependent tasks (green or red squares to indicate correct or incorrect responses, respectively). At the end of each run, an overall accuracy score was provided concerning performance on the tasks requiring a button response.

On the second training day, the task switching aspect of the design was introduced. Participants were given six runs of training, with each run composed of one block for each of 11 tasks that required manual responses. As on day 1, the timing for the first four run was self-paced, with the participants allowed to read the instructions at their own pace prior to initiating the 30 s block. For the final two practice runs, the instruction phase was limited to 5 s, thus introducing the protocol that would be used in the scanner. Training on this day only included tasks that required overt responses. On the third training day, the participants practiced all 17 tasks in four 10 minute runs (35 s/task), emulating the protocol to be used in the scanner sessions.

This training program ensured that participants were familiar with the requirements for each task and had considerable experience in switching between tasks. In this manner, we sought to minimize the impact of learning during the scanning sessions. On the third training day, performance was asymptotic, with the participants correct on at least 85% of the trials for all of the tasks (range = 85% to 98%; see Fig S3).

### Eye tracking

Eye-tracking data were recorded on the third training session to obtain an estimate of saccadic eye movements for each of the tasks. An algorithm implemented in the Eyelink toolbox^33^ identified saccadic eye movements as events in which eye velocity briefly exceeded a threshold of 30 deg/s. These data, tabulated as the mean number of eye movements per task, were included as a motor feature in the second-level feature model (see below). Eye-tracking data from two participants in set A and three participants in set B were not obtained due to technical problems. However, since the eye-movement behavior was consistent across participants, we used group-based estimates.

### Scanning sessions

Participants completed four scanning sessions in total, two with set A and two with set B. The first scanning session for each set was conducted within a few days of the final training session (set A: mean=2.0 days, SD=1.6 days; set B: mean=2.2 days, SD=1.7) and the second scanning session was completed no more than 7 days after the first scanning session (set A: mean=3.1 days, SD=2.5; set B: mean=2.7 days, SD=2.3). Each scanning session consisted of eight imaging runs (10 min total duration/run). Each of the 17 tasks was presented once for 35 s in each imaging run, producing two final sets composed of 16 independent measurements per task. The task order was randomized across runs. To reduce order effects within each set, no two tasks were presented in the same order in two different runs. The order within each run, as well as the order of the runs, was kept constant for all of the participants. This procedure was chosen to allow for cross-participant analyses on the time series level (results not presented here). As noted above, when possible, novel stimuli were used in each run to reduce the recall of specific stimulus-response associations.

### Image acquisition

All fMRI data were acquired on a 3T Siemens Prisma at the Centre for Functional and Metabolic Mapping at Western University. Whole-brain functional images were acquired using an EPI sequence with multi-band acceleration (factor 3, interleaved) with an in-plane acceleration (factor 2), developed at the Centre for Magnetic Resonance Research at the University of Minnesota. Imaging parameters were: TR=1 sec, FOV=20.8cm, phase encoding direction was P to A, acquiring 48 slices with in-plane resolution of 2.5 mm x 2.5 mm and 3 mm thickness. GRE field maps were also acquired for distortion correction of the EPI images due to B0 inhomogeneities (TR=.5 s, FOV=24 cm, 46 slices with in-plane resolution of 3 mm x 3 mm x 3 mm). We also acquired on-line physiological recordings of both heart and respiration during each functional run. No participants had to be excluded from either dataset due to excessive motion. For anatomical localization and normalization, a 5 min high-resolution scan of the whole brain was acquired (MPRAGE, FOV=15.6 cm x 24 cm x 24 cm, at 1x1x1 mm voxel size).

### Image preprocessing

Data preprocessing was carried out using tools from SPM 12^34^, Caret^35^, and SUIT^19^, as well as custom-written scripts written in MATLAB 2015b. For all participants, the anatomical image was acquired in the first scanning session and reoriented to align with the Left-Inferior-Posterior (LPI) coordinate frame. Functional data were re-aligned for head motion within each session, and for different head positions across sessions using the 6-parameter rigid body transformation. The mean functional image was then co-registered to the anatomical image, and this transformation was applied to all functional images. No smoothing or anatomical normalization was applied to the functional images.

### General linear model

A general linear model (GLM) was fit to the time series of each voxel separately for each imaging run. The 5 s instruction phase for all tasks was modeled using a single regressor, but not included in later analyses. Each task was modeled using a boxcar regressor of 30 s, or a combination of multiple regressors if the block contained sub-conditions. These regressors could be 2 boxcar regressors of 15 s each (e.g., N-back task where one sub-condition is 0-Back and the second is 2-Back), 3 boxcar regressors of 10 s each (e.g., visual search, display sizes of 4, 8, or 12), or 2 event-related regressors (e.g., Stroop task, where each trial was congruent or incongruent). The rest condition was not modeled explicitly, but rather used as an implicit baseline in the model.

The quality of the GLM in modeling the BOLD response was determined by measuring the consistency of the activation patterns in the cerebellum across runs. This measure indicated that it was advantageous to omit the traditional high-pass filtering operation before the linear model (default operation in SPM). Instead, we opted to rely on the high-dimensional temporal autocorrelation model (FAST option in SPM) to determine the optimal filtering, implemented in the GLM-estimation. The beta-weights from the first-level GLM were univariately pre-whitened by dividing them by the square root of the residual mean square image. To include rest as a task condition in all subsequent analyses, we added a zero as an estimate for the rest condition and then removed the mean for each voxel across all conditions. As such, the beta-weights expressed the amount of activation elicited by each condition relative to the mean of all conditions.

To combine activation estimates across the two tasks sets, the mean of the shared tasks was removed separately for each set. Both sets were then combined, retaining the repeated estimates for the shared task. This resulted in a total of 61 estimates (set A: 29; set B: 32) for the 47 unique conditions. The activation patterns were re-centered by removing the overall mean across all 61 conditions.

### Cerebellar spatial normalization

The spatially unbiased infratentorial template (SUIT) toolbox (v3.2) in SPM 12 was used to isolate the cerebellum from the rest of the brain and to provide a normalization to a spatially unbiased template of the cerebellum. The segmentation procedure was used to create probability maps of gray and white matter, allowing us to separate cerebellar and cortical tissue. The resulting cerebellar isolation mask was hand corrected to ensure that it did not contain any shared voxels between the superior cerebellum and the directly abutting cerebral cortical regions of the inferior temporal and occipital cortex.

The probabilistic maps for the cerebellum were normalized into SUIT space using the diffeomorphic anatomical registration (DARTEL) algorithm^36^. This algorithm deforms the cerebellum to simultaneously fit the probability maps of cerebellar gray and white matter onto the SUIT atlas template. This non-linear deformation was applied to both the anatomical and functional data. The activation estimates (i.e., the beta weights), and residual mean-square images from the first-level GLM were resliced into SUIT space. All images were masked with the cerebellar mask to avoid activation influences from the inferior occipital cortex. All data were visualized on a surface-based, flat-map representation of the cerebellar cortex in the SUIT toolbox. The flat-map representation allows the spatial extent of task-evoked activation patterns to be fully visualized.

### Motor feature model

Our primary goal was to study the task-evoked activation patterns in the cerebellum beyond the well-known domain of motor function. Although not designed explicitly to measure motor-related activation, the 61 task conditions differed in the number of manual responses, as well as eye movements. To account for these motor-related activations, we generated a motor feature model (task conditions x 3 motor features). For the hand movements, we entered the number of left and right hand presses for each task during the scanner runs (range 1 to 12, combining over the two hands). For eye movements, we used the group-averaged eye movement data from the third training day to estimate the number of saccades for each task condition. All motor features were encoded in terms of movements/s and z-normalized.

To extract and remove the motor-related activation across tasks, the three motor features were combined with an indicator matrix that had a 1 for each of the 61 task conditions, except for rest. To estimate and subsequently remove the influence of the motor features, we estimated the linear model with L2-norm regression (fixed lambda of .01) from the beta-estimates of each participant (task conditions x voxels x participant). The average of all task conditions was used as a baseline measure and subtracted from the motor-corrected activation estimates. The activation estimates for the shared tasks were first averaged and then a group average was computed for the purposes of visualization on the cerebellar flat-map^10^.

### Reliability of activation patterns

To determine intra-subject reliability across the entire cerebellar cortex, we calculated the correlation between the average activation estimates for the first and second session for each task set, separately for each participant. To obtain an overall reliability, we stacked the 29 (A) or 32 (B) activation estimates for all cerebellar voxels into a single vector and calculated the Pearson correlation between the two estimates. For Fig 1e this analysis was also performed for each voxel separately. The group-averaged correlations were then visualized on the cerebellar surface.

### Spatial frequency of activation patterns

To determine how much of the variance of the activation patterns was common to the group relative to how much was idiosyncratic to the individual participants, we calculated two correlations, one between task-activity maps between two sessions for the same participant (as for the reliability), and the second between sessions of different participants. Correlations were computed on all gray matter voxels in SUIT space. To determine the spatial scale of these common activation patterns, we decomposed the volume image for each task condition into five spatial frequency bands ranging from 0 to 5 cycles/cm. This decomposition was done separately for each participant, study, session, and task condition. The within-and between-subject correlations were then computed for each spatial frequency band.

### Evaluating functional boundaries

We developed a novel method to evaluate functional boundaries from fMRI data. The rationale of the method is that, if a boundary is dividing two functionally heterogeneous regions, then two voxels that lie within the same region should have more similar functional profiles than two voxels that are in different regions (Fig 2a; Eq. 1). Because functional organization tends to be smooth, the correlation between two voxels will be higher for two adjacent voxels, and fall off as the spatial distance increases^11^. To control for distance, we calculated the activation pattern correlations for all pairs of voxels separated by a fixed Euclidean distance, using spatial bins ranging from 4 mm to 35 mm. This analysis was conducted in the volume, as a veridical surface reconstruction of the cerebellar folia is only possible for resolutions better than 0.2mm^37^. This should slightly favor parcellations that align better with lobular boundaries, as two voxels on opposite sides of a fissure tend to be further apart on the cerebellar cortex than two voxels on the same side, even if their distance in the volume is matched. To exclude spatial correlations that were driven by noise, we used a cross-validated correlation. Using **u**_i,1_ to represent the functional profile (zero-meaned) of voxel i from one session, and **u**_j,2_ the functional profile of voxel j from the other session, the correlation was calculated as:

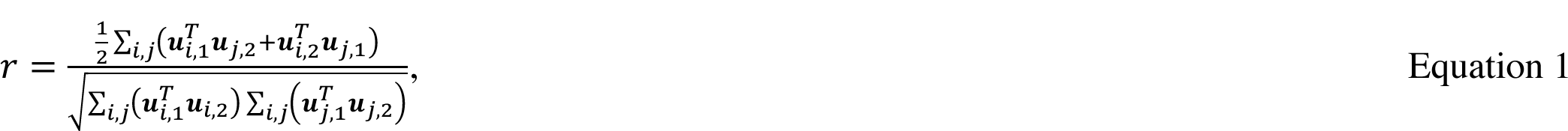

where the sum was done on all voxel pairs *i,j* in the corresponding 5 mm bin, separating those that involved voxel pairs from the same region (within) and those in which the voxel pairs spanned the boundary (between) for each spatial bin. We excluded voxels where the term 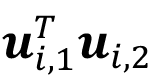 was negative, as it indicated the absence of any reliable tuning across the two sessions. The difference between the within-region correlation and the between-region correlation defined the distance controlled boundary coefficient (DCBC). A positive DCBC value indicates that voxel pairs originating from the same region are more functionally related than voxel pairs that lie across boundaries. The DCBC was calculated for each participant and spatial bin separately, and then averaged.

The DCBC can serve not only as a global measure of a parcellation (averaging across the cerebellum and spatial bins), but also as a measure to evaluate the strength of individual boundaries. For the latter, we first identified boundaries using an edge-based connectivity scheme^79^. The strength of a given boundary is defined by the DCBC calculated only on the voxel pairs from the two regions that are separated by that boundary. To visualize boundary strength, the thickness of the boundary on the flat-map was based on its DCBC value.

We applied this boundary evaluation procedure to MDTB parcellations, as well as parcellations based on lobular boundaries or task-free fMRI data. The lobular parcellation was obtained from a probabilistic atlas of the human cerebellum^19^ that includes lobules I-IV, V, VI, Crus I, Crus II, VIIb, VIIIa, VIIIa, IX, and X. To ensure that that poor performance of the lobular parcellation was not due to inaccuracies in detecting the lobular boundaries, we repeated the analysis using a manual lobular parcellation in five participants. The parcellation from this sample predicted functional boundaries about as well as the one derived from the probabilistic atlas (DCBC: .022 vs. .025; t_5_=1.441. p=0.209). The task-free 10-region parcellation^6^ was based on data archived as part of the Human Connectome Project (HCP), while the other two were based on a large 1000-person dataset collected at Harvard and Massachusetts General Hospital^5^.All parcellations were sampled into SUIT space and evaluated using our multi-domain task battery.

### Multi-domain task dataset (MDTB) parcellation

To derive a parcellation from the MDTB, we used the activation profile averaged in SUIT volumetric space across participants. The parcellation was based only on data from voxels assigned to gray matter. As a clustering approach, we used convex nonnegative matrix factorization^15^, which decomposes the N (tasks) x P (voxels) data matrix into a product of an N x Q (regions) matrix of task profiles and an Q x P matrix of voxel weights. The voxel weights, but not the task profiles, are constrained to be non-negative. Furthermore, the task profiles are convex combinations of the raw data. In comparison to other decomposition methods, such as independent component analysis (ICA), this method has the advantage that voxels cannot be explained by an inverted or negative regional task profile. This constraint is also reasonable given that, the main neural signal driving the BOLD response in the cerebellum, the mossy fiber input, is excitatory^27^. A winner-take-all approach was adopted to assign each voxel to the region with the highest weight. To ensure convergence, we started the decomposition with random initializations, and selected the iteration with the best reconstruction of the original data. We stopped when the current estimate of the best solution was obtained five times without being replaced by a better solution. The results from this winner-take-all approach were then graphically displayed on a flat-map of the cerebellum.

To allow for a direct comparison with existing task-free parcellations^5, 6^, we used parcellations with 7, 10, and 17 regions (Fig S5d-f). Parcellations involving regions within this range achieved similar reconstruction accuracy and quality of functional boundaries.

We also derived separate parcellations for each participant to determine whether boundaries could be predicted on the individual. An advantage of the individual parcellation is that the idiosyncrasies of within-subject variability is captured, which of course is missed at the group level. The disadvantage is that individual parcellations are derived on substantially less data than the group.

#### Bootstrap analysis

To obtain a measure of boundary uncertainty, we performed bootstrap analyses across both participants and task conditions. For the participant bootstrap, we repeatedly drew 24 participants (with replacement) from our sample, averaged the data, and derived a new functional parcellation. To be able to relate the parcellations to each other, each parcellation used the original solution as a starting value. For the task bootstrap, we repeatedly drew 47 task conditions (with replacement) from our data, again deriving a new parcellation each time. For each analysis, we repeated this process 100 times.

To evaluate the consistency of the parcellations globally, we calculated the adjusted Rand Index, which measures the correspondence between two parcellations (0: overlap not different from chance, 1: perfect overlap). For a regionally specific analysis, we counted the number of times that each voxel was assigned to the same (most frequent) region. For visual display, we then used this assignment certainty to determine the transparency of the region coloring (<50%: fully transparent, 100%: fully opaque).

#### Evaluation of functional parcellations

To evaluate the group and individual MDTB parcellations, we wanted to know how well functional boundaries could be predicted for each participant, using a completely novel set of tasks. Because we did not acquire data with a third, independent task set on the same participants, we used the existing data to estimate lower and upper bounds of predictability. For the lower bound, we derived the parcellation using the data from set A and evaluated it with the data from set B, using the unique tasks only. This procedure was then repeated with the task sets reversed, and the results were averaged across the two cross-validation folds. Note that the outcome of this analysis will likely result in a lower value than would be obtained with the final segmentation, as each parcellation is based on half of the available data. As such, we use this estimate as an approximate lower bound. We also evaluated a parcellation derived from both sets A and B. We evaluated this parcellation as before, excluding the shared tasks from both task sets to make the estimate consistent with the lower-bound estimate. Because there is overlap between the data used for training and evaluation, the performance measure here is overfit and, therefore, was taken as an approximate upper bound. In sum, we assume that true performance of the full parcellation, if applied to a completely new task set, would fall somewhere between these lower and upper bounds.

The anatomical and task-free parcellations could be directly evaluated on the data from sets A and B since each parcellation was derived from independent data. For consistency, we excluded the data from the shared tasks in the evaluation set.

To evaluate the degree to which the results depended on the similarity of the tasks in the training and test sets, we repeated the analysis, this time selecting the 7 most distinct task conditions in each test set (Fig S6). The conditions were selected by computing the distance between activity patterns for each test condition to each training condition (Fig S4a). We then identified, for each test condition the closest match in the training set and selected the 7 test conditions for which this closest match was most dissimilar.

To further validate our results, we evaluated the MDTB and task-free parcellations on the task-based data from the HCP dataset (https://db.humanconnectome.org). We utilized data from the 214 most recently added participants (scanned at 3T). Of the 214, 186 participants had complete data sets, as these constituted our final sample. For each participant, we evaluated the parcellations on a set of 22 contrast maps from 7 tasks (all against rest).

### Representational structure of task-related activation patterns

Representational similarity analysis^16^ (RSA) was used to investigate the representational structure of task-related activation patterns from the MDTB cerebellar data. The dissimilarity between the motor-corrected activation patterns was measured for each pair of task conditions using the cross-validated Mahalanobis distance, using the imaging runs as independent partitions. To calculate the distances between conditions across the sets, we subtracted the mean of the shared task conditions from each imaging run first. Cross-validation ensures that the average (expected) value of the dissimilarity measure is zero if the two activation patterns only differ by noise. This allowed us to test for significant differences between activation patterns using a one-sample t-test against zero.

Classical multidimensional scaling (MDS) was employed to visualize the distances between all possible pairs of task conditions. For the purposes of visualization, the pairwise distances for the shared tasks were averaged so that there were 47 (rather than 61) task conditions in the representational dissimilarity matrix (RDM). MDS projects the *N*-dimensional RDM into a lower-dimensional space so that distances from the higher space are preserved with as much integrity as possible. MDS was performed on the group-averaged RDM, and the first three dimensions were visualized in a 3-dimensional space.

### Feature-based approach

The power of our task-based approach in studying the cerebellum is that we can identify the involvement of each region across functional domains and different task variations. To summarize the task activation profiles for each region, we used a feature-based encoding method. The features included the three motor features (see above) and cognitive features, selected to capture the hypothetical mental processes involved in each task. To derive these features, we used an online cognitive ontology^17^, an atlas of tasks and the concepts associated with those tasks. Of the 815 concepts currently included in the atlas, 46 were judged to provide an appropriate and sufficient characterization of the tasks in our battery, creating a feature matrix (47 task conditions x 46 features). For example, features such as semantic knowledge and lexical processing were associated with tasks such as verb generation and semantic prediction; emotion recognition was associated with the IAPS emotional processing task and the biological motion task. As with the motor feature model, each feature was z-standardized and feature weights for each region were estimated with non-negative regression. For visualization purposes, the three highest weights for each region were computed.

## Data availability

The activation maps and functional parcellations are made available on diedrichsenlab.org/imaging/mdtb.htm. The raw behavioral and imaging data for the cerebellum is also available on the data-sharing repository openneuro.org. Experimental code is available at github.com/maedbhk/MDTB-Cerebellum.

## Supplementary Figures

**Table S1.** Task set description for all 26 unique tasks and 47 unique conditions. Tasks that require overt motor responses are executed either with the left, right, or both hands.

| Task Name | Task Description | Dataset | Conditions | Hand Assignment |
| --- | --- | --- | --- | --- |
| Objects | Passive viewing, pictures of objects and a checkerboard pattern. | A |  | None |
| Motor Imagery<sup>a</sup> | Imagine playing a game of tennis. | A |  | None |
| Stroop<sup>a</sup> | 3AFC, indicating color of stimulus word (3 colors), comparing conditions in which color-word mapping is congruent or incongruent (Stroop task). | A | Congruent<br>Incongruent | Both |
| Verbal Working Memory<sup>a</sup> | 2AFC, indicating if current stimulus in stream of letters matches letter displayed two items previously (2-back). | A | 2-Back<br>0-Back | Left |
| Interval Timing<sup>a</sup> | 2AFC, indicating if a tone is short (100ms) or long (175ms) | A |  | Right |
| Arithmetic<sup>a</sup> | 2AFC, indicating if simple multiplication equations (e.g. $2 \times 7 = 14$) are correct or incorrect. For control task, participants view a series of four numbers and indicate presence/absence of target number (e.g., 1). | A | Math<br>Digit Judgment | Right |
| IAPS affective<sup>a</sup> | 2AFC, indicating if picture (scenes, animals, foods) is pleasant or unpleasant. | A | Pleasant<br>Unpleasant | Left |
| IAPS emotion | 2AFC, indicating if picture depicts sad or happy face. | A | Happy<br>Sad | Right |
| Go/No-Go<sup>a</sup> | Go-NoGo task with positive (Go) or negative (No Go) words. | A | Go<br>No Go | Left |
| Theory of Mind<sup>a</sup> | 2AFC to indicate if short story contains true or false belief (Theory of Mind task) | A & B |  | Left |
| Rest | Passive viewing of fixation cross. | A & B |  | None |
| Object N-Back<sup>a</sup> | As above, with objects instead of letters (2-back). | A & B | 2-Back<br>0-Back | Right |
| Verb Generation<sup>a</sup> | Verb generation task requiring covert responses to visually-presented nouns, either repeating the stimulus (Read) or generating a verb associated with the noun (Generate). | A & B | Verb Generation<br>Word Reading | None |
| Spatial Imagery<sup>a</sup> | Imagine walking from room to room in childhood home, with a cue specifying the path to be taken (e.g., "Imagine walking from the kitchen to the bedroom, stopping to look around at different rooms"). | A & B |  | None |
| Motor Sequence<sup>a</sup> | 6-element sequence, either requiring one key press with each of six fingers (bimanual) or repetition of a single key press with one finger (unimanual left or right). | A & B | Finger Sequence<br>Finger Simple | Both |
| Action Observation<sup>a</sup> | Passive viewing of videos of knots being tied, learning the name of the knot (presented at top of screen) for a latter recall test. | A & B | Action Observation<br>Video Knots | None |
| Visual Search<sup>a</sup> | 2AFC, indicating if target stimulus (“L”) is present among distractors (“T”), with varying set size (4, 8, 12). | A & B | Easy (4)<br>Medium (8)<br>Hard (12) | Left |
| Spatial Map | Memorize a spatial mapping of numbers (either, 1, 4, or 7) for subsequent recall | B | Easy (1)<br>Medium (4)<br>Hard (7) | Both |
| Mental Rotation<sup>a</sup> | Mentally rotate target object to determine whether it can be brought into alignment with baseline object. Difficulty is measured by angular disparity between target and baseline image. Stimuli were obtained from Ganis and Kievit (2015)<sup>a</sup> | B | Easy (0)<br>Medium (50)<br>Difficult (150) | Right |
| Biological Motion<sup>a</sup> | 2AFC to identify intact point-light walkers (either happy or sad) or scrambled walkers (fast or slow). Stimuli obtained from Troje et al. (2017)<sup>a</sup> | B | Biological Motion<br>Scrambled Motion | Right |
| Concrete Permuted Rules Operations (CPRO)<sup>a</sup> | Apply task-rule set (logic, sensory, & motor rules) to two consecutively presented stimuli (rectangles: either red or blue, vertical or horizontal) | B |  | Both |
| Word Prediction<sup>a</sup> | 2AFC task to indicate if five sequentially-presented words comprise a semantically meaningful sentence. Stimuli obtained from D’Mello et al. (2017)<sup>a</sup> | B | True Prediction<br>Violated Prediction<br>Scrambled Prediction | Left |
| Response Alternatives<sup>a</sup> | Execute a fast motor response to an imperative signal (white cross) that appears in one of 1, 2, or 4 primed positions | B | Easy (1)<br>Medium (2)<br>Hard (4) | Both |
| Nature Movie<sup>a</sup> | Passive viewing of a nature clip of kickboxing kangaroos, taken from “Planet Earth II: Islands” | B |  | None |
| Animated Movie<sup>a</sup> | Passive viewing of an emotional love story between two characters from the Pixar movie “Up” | B |  | None |
| Landscape Movie<sup>a</sup> | Passive viewing of an aesthetically-pleasing clip that depicts a diverse scenery, taken from Vimeo | B |  | None |

**Figure S1.**
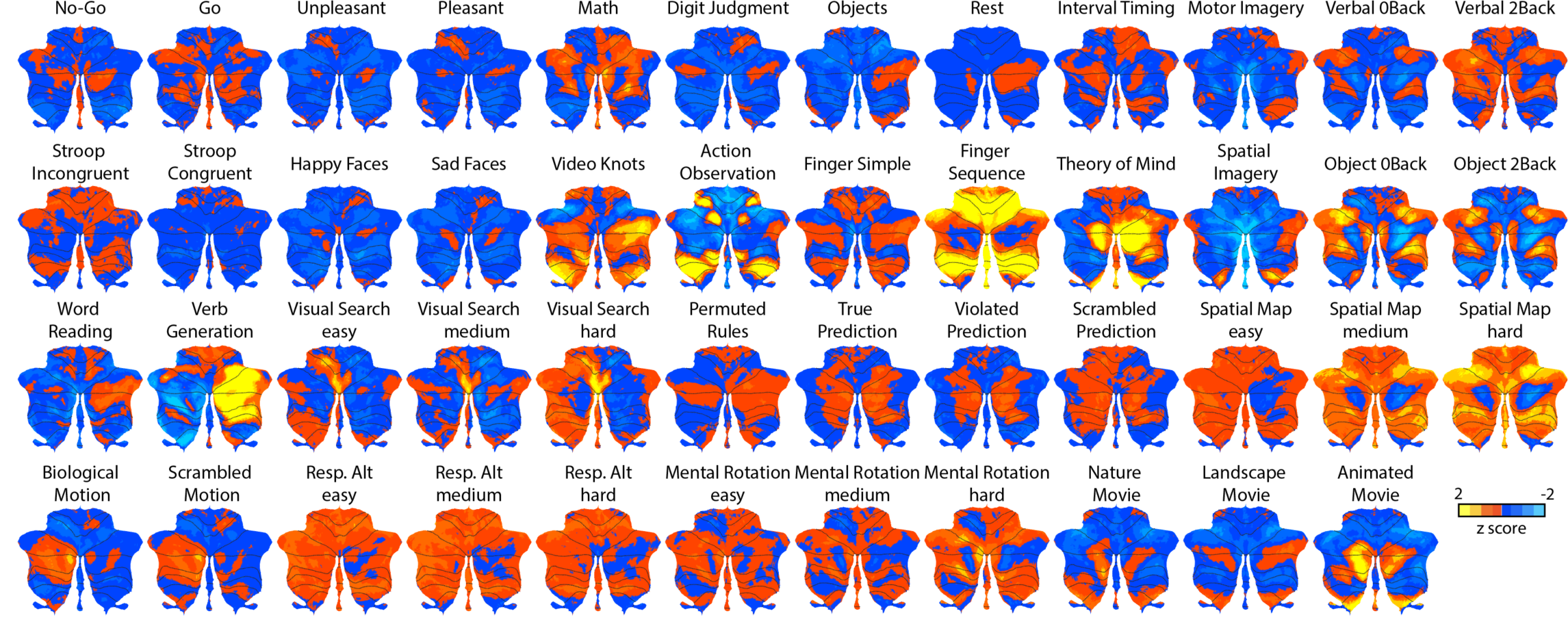
Unthresholded, group-averaged activation maps for the 47 unique task conditions displayed on a surface-based representation of the cerebellar cortex^10^. All activations are calculated relative to the mean activation across all conditions. Red-to-yellow colors indicate increases in activation and blue colors indicate decreases in activation.

**Figure S2.**
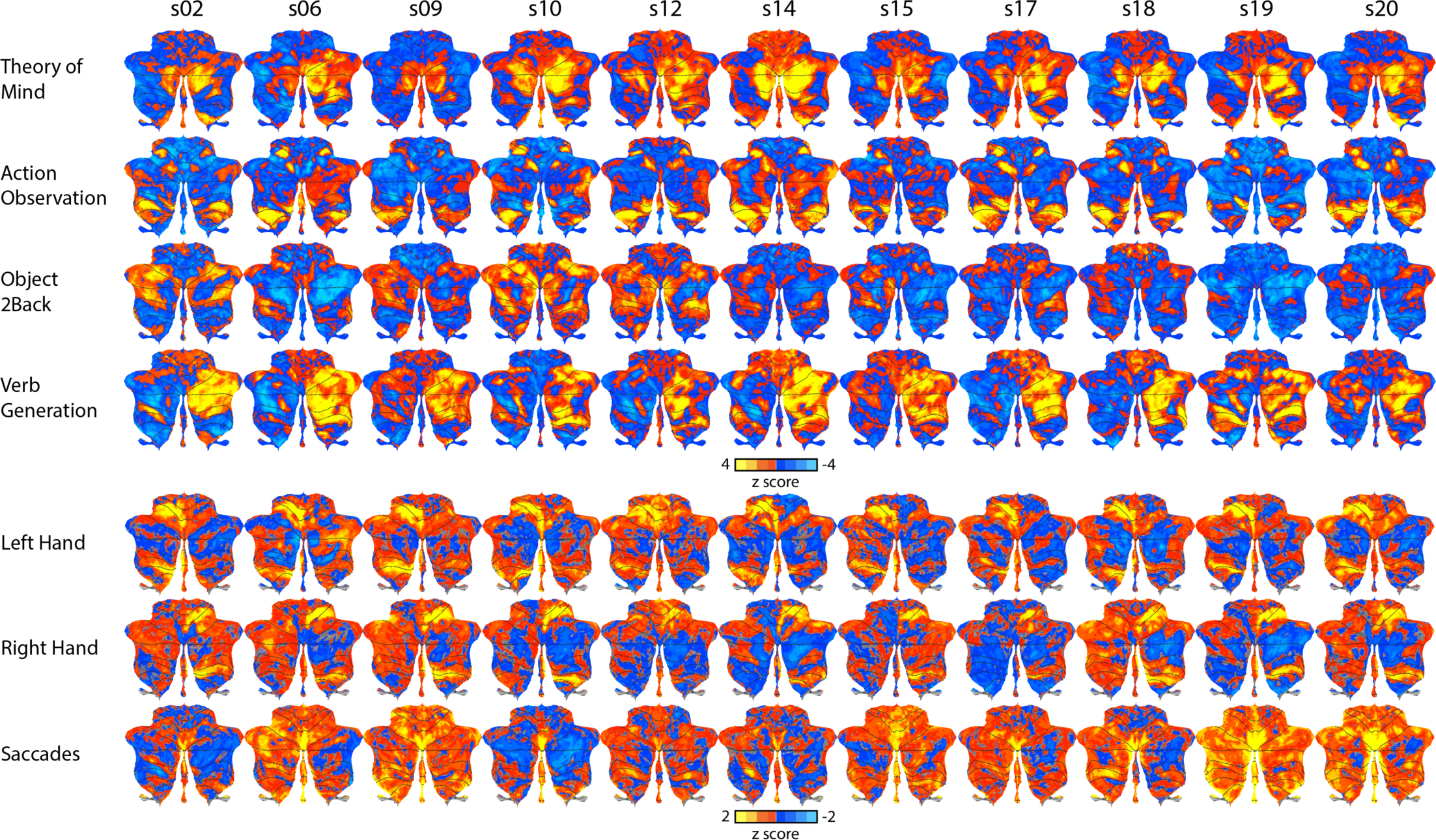
Unthresholded, individual activation maps for 4 representative tasks and motor feature maps for 11 representative participants. All activations are calculated relative to the mean activation across all conditions. Red-to-yellow colors indicate increases in activation and blue colors indicate decreases in activation.

**Figure S3.**
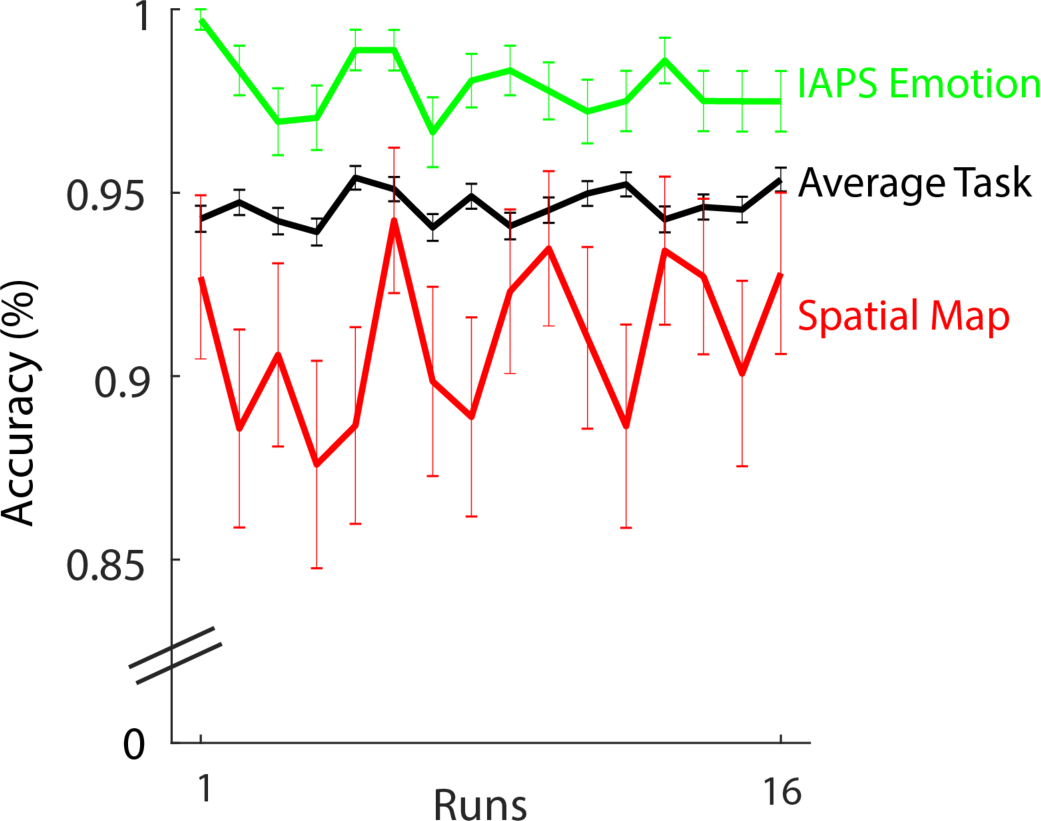
Task performance (% accuracy) averaged across the four scanning sessions, each composed of four runs. Average across all tasks is shown in black. Poorest performance was on the spatial map task (red line) and best performance was on the IAPS emotion task (green line).

**Figure S4.**
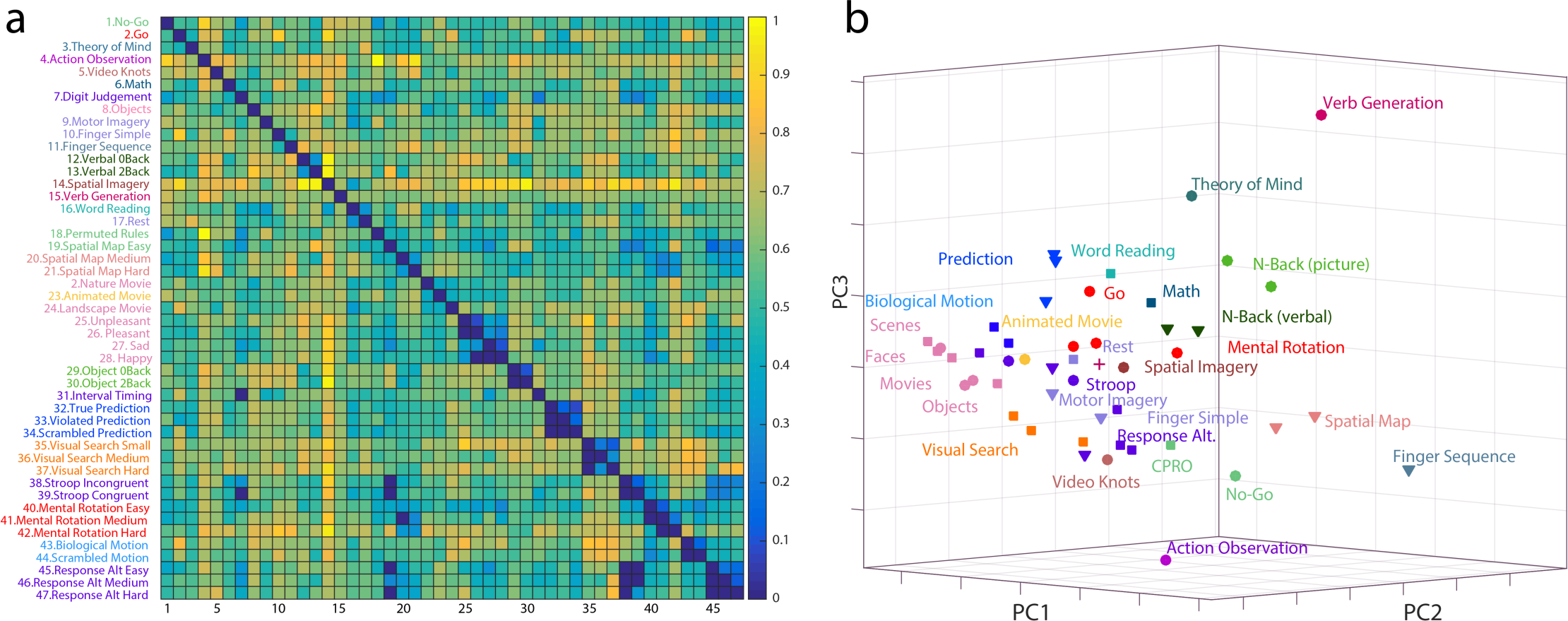
Representational task space for 47 unique task conditions presented in a representational dissimilarity matrix (RDM) and a multi-dimensional scaling (MDS). **(a)** RDM for the group-averaged data for the unique 47 task conditions. Shared tasks are averaged across the four scanning sessions. Dark blue represents low dissimilarity between pairwise task-evoked activity patterns while high distances (bright yellow) represent high dissimilarity between pairwise task-evoked activity patterns. Thresholded values are shown below the diagonal (dark blue cells indicating pairwise comparisons between task conditions were not significant (p<.001, e.g., pleasant and unpleasant scenes). **(b)** A multi-dimensional scaling plot (using first three PCs for display purposes), showing the relative similarity of the task-evoked activity patterns after correction for activity related to basic motor output. Hierarchical clustering is applied to the tasks, with colors in both the RDM and MDS indicating cluster membership.

**Figure S5.**
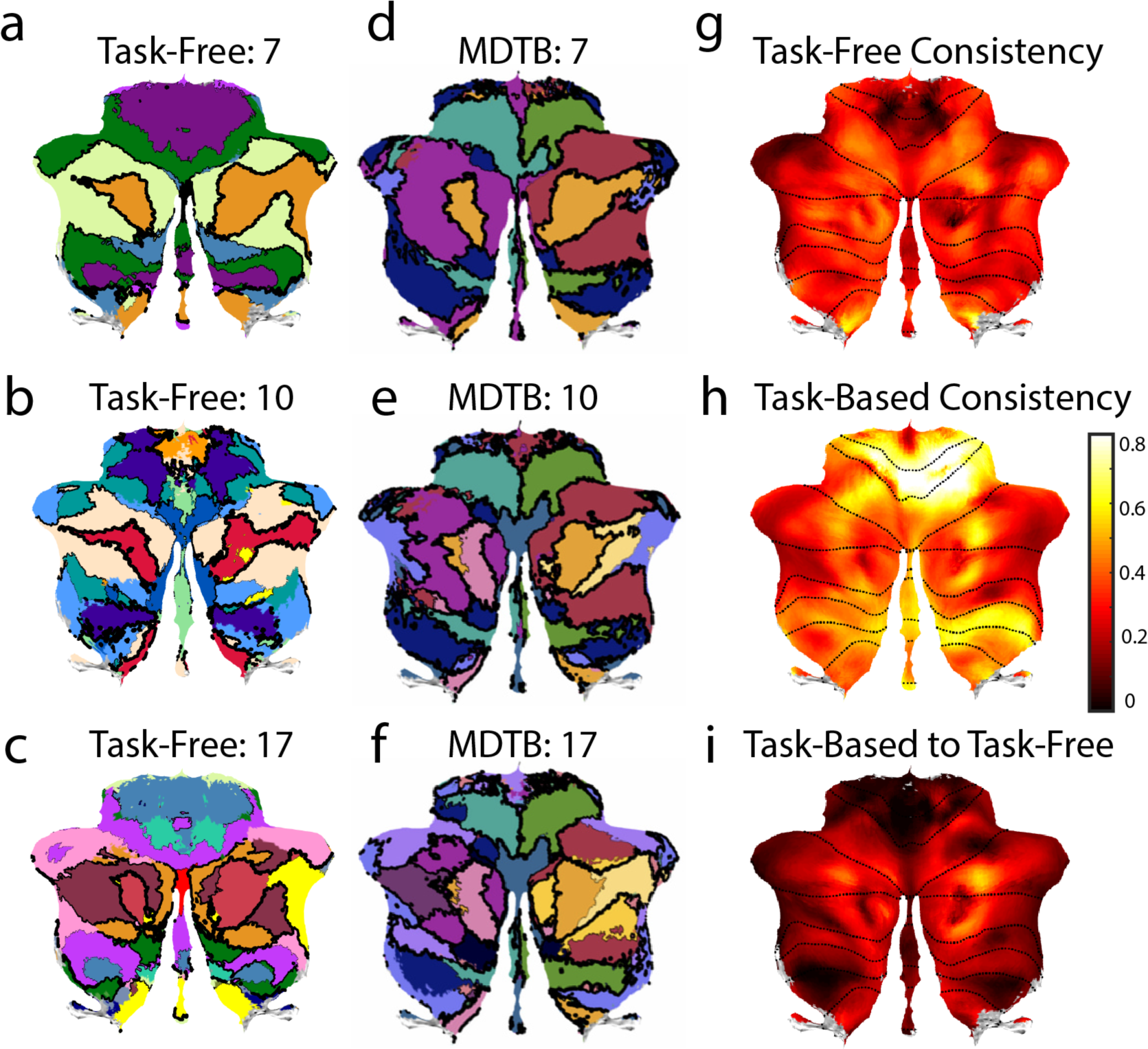
7, 10, and 17 region parcellations derived from task-free HCP **(a-c)** and MDTB **(d-f)** data. **(g**) Average Rand coefficient between task-free parcellations, computed locally (1cm sphere) around each cerebellar voxel. **(h)** Average Rand coefficient between MDTB parcellations. **(i)** Average difference of Rand coefficients for the MDTB and task-free parcellations.

**Figure S6.**
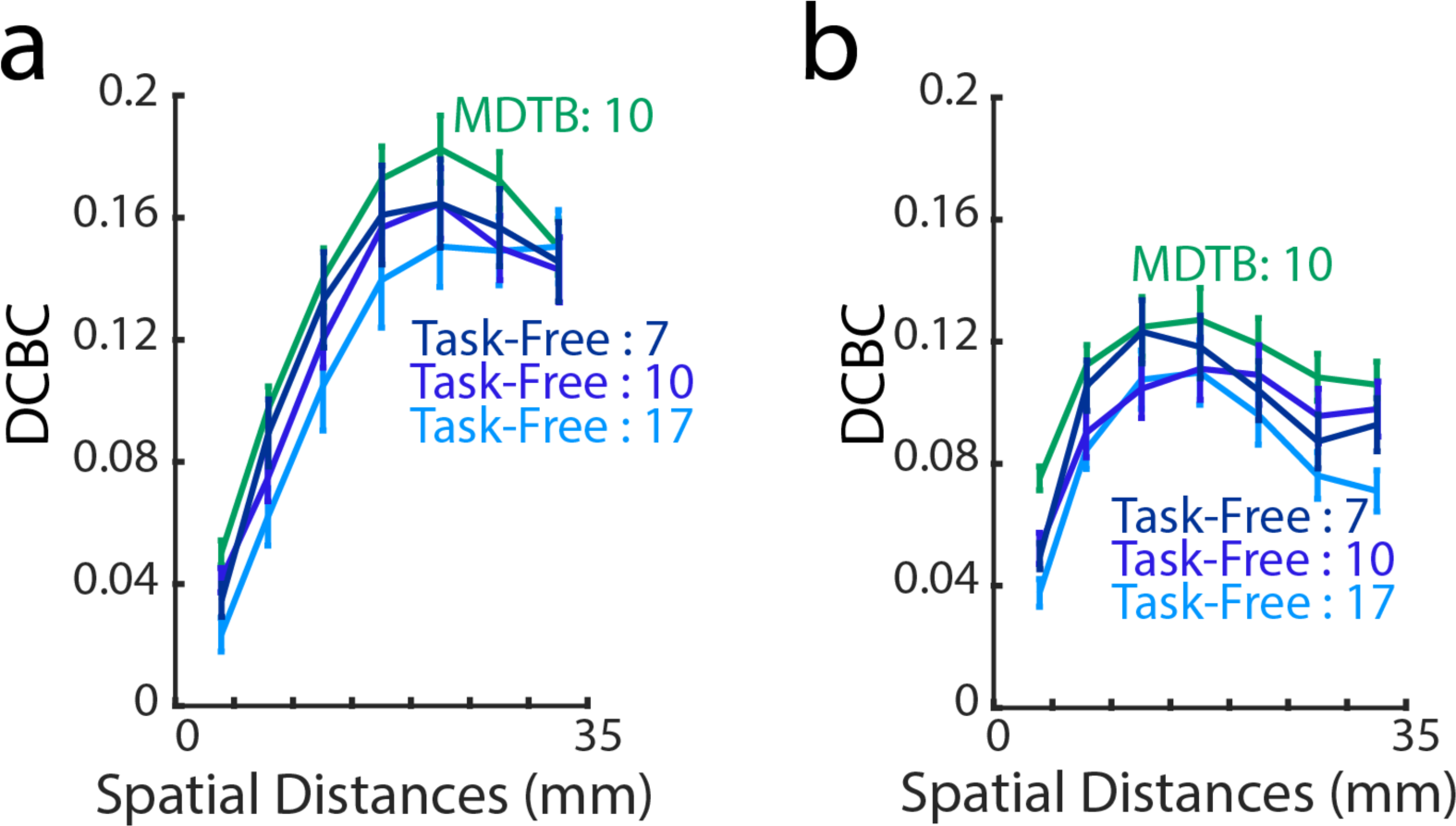
Cross-validated evaluation of MDTB parcellation on a subset of 7 tasks, selected to be most dissimilar to task conditions included in the data set. For comparison purposes, task-free parcellations are evaluated on the same tasks. **(a)** MDTB parcellation trained on Set A and evaluated on 7 tasks from Set B (Mental Rotation Easy, Mental Rotation Medium, Mental Rotation Hard, Spatial Map Medium, Spatial Map Hard, Animated Movie, and Nature Movie). **(b)** MDTB parcellation trained on Set B and evaluated on 7 tasks from Set A (Sad Faces, Interval Timing, Go, Theory of Mind, Word Reading, Motor Imagery, Math).

**Figure S7.**
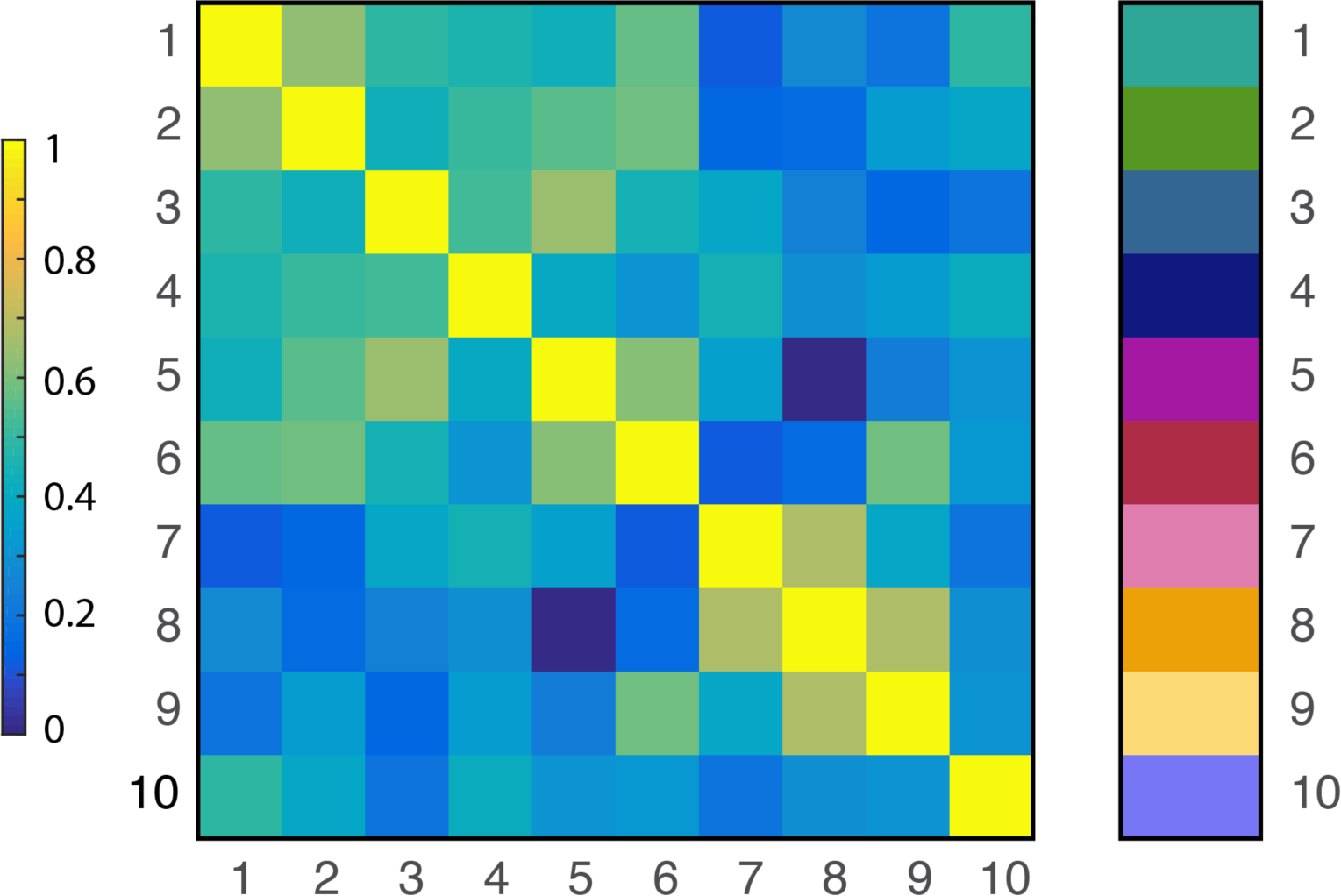
Correlation between the task-profiles of the 10 regions of the MDTB parcellation. The values in the correlation matrix are scaled between 0 (blue) and 1 (yellow). The bar on the right denotes the colors of each of the 10 regions (see Fig 5).

## Acknowledgements

This work was supported by a James S. McDonnell Foundation (Scholar award to J.D.), the Canadian Institutes of Health Research (PJT 159520 to J.D.), a Platform Support Grant from Brain Canada and the Canada First Research Excellence Fund (BrainsCAN) to Western University, the National Science Foundation (OAC-1649658 to R.P.), and the National Institute of Health (NS092079, NS105839 to R.I.). Data from the Human Connectome Project were analyzed, WU-Minn Consortium (Principal Investigators: David Van Essen and Kamil Ugurbil; 1U54MH091657) funded by the 16 NIH Institutes and Centers that support the NIH Blueprint for Neuroscience Research, and by the McDonnell Center for Systems Neuroscience at Washington University. Thanks to Jon Walters for assistance in task annotation.

## Contributions

R.I. and J.D. originally conceived of the project. M.K., J.D., and R.I. designed the study. M.K. and C.H. collected the data. M.K. and J.D. performed the analyses. R.P., M.K., J.D., and R.I. annotated the cognitive tasks. M.K., J.D., and R.I. wrote the manuscript. All authors discussed the results and contributed to the final manuscript.

## References

1. Ivry, R. B. & Baldo, J. V. Is the cerebellum involved in learning and cognition? Curr. Opin. Neurobiol. 2, 212–216 (1992).

2. Kelly, R. M. & Strick, P. L. Cerebellar loops with motor cortex and prefrontal cortex of a nonhuman primate. J. Neurosci. 23, 8432–8444 (2003).

3. Allen, G., Buxton, R. B., Wong, E. C. & Courchesne, E. Attentional activation of the cerebellum independent of motor involvement. Science (80-.). 275, 1940–1943 (1997).

4. Larsell, O. The development of the cerebellum in man in relation to its comparative anatomy. J. Comp. Neurol. (1947).

5. Buckner, R. L., Krienen, F. M., Castellanos, A., Diaz, J. C. & Yeo, B. T. T. The organization of the human cerebellum estimated by intrinsic functional connectivity. J Neurophysiol 106, 2322– 2345 (2011).

6. Spronk, Kulkarni, Repvos, Anticevic & Cole, M. W. Mapping the human brain’s cortical-subcortical functional network organization. (2017).

7. Guell, X., Schmahmann, J. D., Gabrieli, J. D. E. & Ghosh, S. S. Functional Gradients of the Cerebellum: a Fundamental Movement-to-Thought Principle. 1–43 (2018).

8. Stoodley, C. J., Valera, E. M. & Schmahmann, J. D. Functional topography of the cerebellum for motor and cognitive tasks: An fMRI study. Neuroimage 59, 1560–1570 (2012).

9. Stoodley, C. J. & Schmahmann, J. D. Functional topography in the human cerebellum: A meta-analysis of neuroimaging studies. Neuroimage 44, 489–501 (2009).

10. Diedrichsen, J. & Zotow, E. Surface-based display of volume-averaged cerebellar imaging data. PLoS One 10, 1–18 (2015).

11. Wiestler, T., Mcgonigle, D. J. & Diedrichsen, J. Integration of sensory and motor representations of single fingers in the human cerebellum. 3042–3053 (2011). doi:10.1152/jn.00106.2011.

12. Ohtsuka, K. & Noda, H. Discharge Properties of Purkinje-Cells in the Oculomotor Vermis During Visually Guided Saccades in the Macaque Monkey. J. Neurophysiol. 74, 1828–1840 (1995).

13. Nitschke, M. F., et al. Activation of Cerebellar Hemispheres in Spatial Memorization of Saccadic Eye Movements : An fMRI Study. 164, 155–164 (2004).

14. Horak, F. B. & Diener, H. C. Cerebellar control of postural scaling and central set in stance. J Neurophysiol 72, 479–493 (1994).

15. Ding, C., Li, T. & Jordan, M. I. Convex and Semi-Nonnegative Matrix Factorizations. 1–26 (2008).

16. Diedrichsen, J. & Kriegeskorte, N. *Representational models: A common framework for understanding encoding, pattern-component, and representational-similarity analysis*. PLoS Computational Biology 13, (2017).

17. Poldrack, R. A. et al. The Cognitive Atlas: Toward a Knowledge Foundation for Cognitive Neuroscience. Front. Neuroinform. 5, 1–11 (2011).

18. Marek, S. et al. Spatial and Temporal Organization of the Individual Human Cerebellum. SSRN Electron. J. 1–17 (2018). doi:10.2139/ssrn.3188429

19. Diedrichsen, J., Balsters, J. H., Flavell, J., Cussans, E. & Ramnani, N. A probabilistic MR atlas of the human cerebellum. Neuroimage 46, 39–46 (2009).

20. Airey, D. C., Lu, L. & Williams, R. W. Genetic control of the mouse cerebellum: Identification of quantitative trait loci modulating size and architecture. J. Neurosci. 21, 5099–5109 (2001).

21. Apps, R. & Hawkes, R. Cerebellar cortical organization: A one-map hypothesis. Nat. Rev. Neurosci. 10, 670–681 (2009).

22. Buckner, R. L., Krienen, F. M., Castellanos, A., Diaz, J. C. & Yeo, B. T. T. The organization of the human cerebellum estimated by intrinsic functional connectivity. J Neurophysiol 106, 2322– 2345 (2011).

23. Buckner, R. L., Krienen, F. M., Castellanos, A., Diaz, J. C. & Yeo, B. T. T. The organization of the human cerebellum estimated by intrinsic functional connectivity. J Neurophysiol 106, 2322– 2345 (2011).

24. Guell, X., Gabrieli, J. D. E. & Schmahmann, J. D. Triple representation of language, working memory, social and emotion processing in the cerebellum: convergent evidence from task and seed-based resting-state fMRI analyses in a single large cohort. Neuroimage 172, 437–449 (2018).

25. Sugihara, I. & Shinoda, Y. Molecular, Topographic, and Functional Organization of the Cerebellar Cortex: A Study with Combined Aldolase C and Olivocerebellar Labeling. J. Neurosci. 24, 8771–8785 (2004).

26. Leclerc, N., Doré, L., Parent, A. & Hawkes, R. The compartmentalization of the monkey and rat cerebellar cortex: zebrin I and cytochrome oxidase. Brain Res. 506, 70–78 (1990).

27. Lauritzen, M. Relationship of spikes, synaptic activity, and local changes of cerebral blood flow. J Cereb Blood Flow Metab 21, 1367–1383 (2001).

28. Hawkes, R., Gallagher, E. & Ozol, K. Blebs in the mouse cerebellar granular layer as a sign of structural inhomogeneity 1. Anterior lobe vermis. Cells Tissues Organs 158, 205–214 (1997).

29. Tavor, I. et al. Task-free MRI predicts individual differences in brain activity during task performance. Science 352, 216–20 (2016).

30. Martin, T. A., Keating, J. G., Goodkin, H. P., Bastian, A. J. & Thach, W. T. Throwing while looking through prisms I. Focal olivocerebellar lesions impair adaptation. Brain 119, 1183–1198 (1996).

31. Andreasen, N. C. & Pierson, R. The Role of the Cerebellum in Schizophrenia. Biol. Psychiatry 64, 81–88 (2008).

32. Moberget, T. et al. Cerebellar volume and cerebellocerebral structural covariance in schizophrenia: a multisite mega-analysis of 983 patients and 1349 healthy controls. Mol. Psychiatry 23, 1512–1520 (2017).

33. Cornelissen, F. W. & Peters, E. M. The Eyelink Toolbox : Eye tracking with MATLAB and the Psychophysics Toolbox. 34, 613–617 (2002).

34. Ashburner, J., Chen, C., Moran, R., Henson, R. & Phillips, C. SPM12 Manual The FIL Methods Group (and honorary members). 1–508

35. Essen, D. C. Van. NeuroImage Cortical cartography and Caret software ☆. Neuroimage 62, 757–764 (2012).

36. Ashburner, J. A fast diffeomorphic image registration algorithm. 38, 95–113 (2007).

37. Sereno, M. I., Diedrichsen, J., Tachrout, M., Silva, G. & De Zeeuw, C. Reconstruction and unfolding of the human cerebellar cortex from high-resolution post-mortem MRI. in Society for Neuroscience (SfN) (2014).

38. Boly, M., et al. When thoughts become action : An fMRI paradigm to study volitional brain activity in non-communicative brain injured patients. Hum. Brain Mapp. J. 36, 979–992 (2007).

39. Egner, T. & Hirsch, J. The neural correlates and functional integration of cognitive control in a Stroop task. 24, 539–547 (2005).

40. Owen, A. M., McMillan, K. M., Laird, A. R. & Bullmore, E. N-back working memory paradigm: A meta-analysis of normative functional neuroimaging studies. Hum. Brain Mapp. 25, 46–59 (2005).

41. Schubotz, R. I. & Cramon, D. Y. Von. Interval and Ordinal Properties of Sequences Are Associated with Distinct Premotor Areas. 210–222 (2001).

42. Rickard, T.. et al. The calculating brain: an fMRI study. Neuropsychologia 38, 325–335 (2000).

43. Moulton, E. A. et al. Aversion-Related Circuitry in the Cerebellum : Responses to Noxious Heat and Unpleasant Images. 31, 3795–3804 (2011).

44. Elliott, R., Rubinsztein, C. A. J. S., Sahakian, B. J. & Dolan, R. J. Selective attention to emotional stimuli in a verbal go / no-go task : an fMRI study. 11, 1739–1744 (2000).

45. Dodell-feder, D., Koster-hale, J., Bedny, M. & Saxe, R. NeuroImage fMRI item analysis in a theory of mind task. Neuroimage 55, 705–712 (2011).

46. Fiez, J. A. Cerebellar Contributinos to Cognition. Neuron 16, 13–15 (1996).

47. Wiestler, T. & Diedrichsen, J. Skill learning strengthens cortical representations of motor sequences. Elife 2013, 1–20 (2013).

48. Cross, E. S. et al. Physical experience leads to enhanced object perception in parietal cortex: Insights from knot tying. Neuropsychologia 50, 3207–3217 (2012).

49. Donner, T. H., et al. Visual Feature and Conjunction Searches of Equal Difficulty Engage Only Partially Overlapping Frontoparietal Networks. 25, 16–25 (2002).

50. Shepard, R. N. & Metzler, J. Mental Rotation of Three-Dimensional Objects. Science (80-.). 171, 701–703 (1971).

51. Ganis, G. & Kievit, R. A. A new set of three-dimensional shapes for investigating mental rotation processes: validation data and stimulus set. J. Chem. Inf. Model. 53, 1689–1699 (2013).

52. Troje, N. F. Decomposing biological motion : A framework for analysis and synthesis of human gait patterns. 371–387 (2017).

53. Ito, A. T. et al. Cognitive task information is transferred between brain regions via resting-state network topology. Nat. Commun. 8, 1–31 (2017).

54. Moberget, T., Gullesen, E. H., Andersson, S., Ivry, R. B. & Endestad, T. Generalized Role for the Cerebellum in Encoding Internal Models: Evidence from Semantic Processing. J. Neurosci. 34, 2871–2878 (2014).

55. D’Mello, A. M., Turkeltaub, P. E. & Stoodley, C. J. Cerebellar tDCS modulates neural circuits during semantic prediction: A combined tDCS-fMRI study. J. Neurosci. (2017). doi:10.1523/JNEUROSCI.2818-16.2017

56. Bischoff-Grethe, A., Ivry, R. B. & Grafton, S. T. Cerebellar involvement in response reassignment rather than attention. J. Neurosci. 22, 546–553 (2002).

57. Nguyen, V. T., et al. Distinct Cerebellar Contributions to Cognitive-Perceptual Dynamics During Natural Viewing. 1–11 (2017). doi:10.1093/cercor/bhw334

